# Spatial transcriptomics reveals BMP-dependent stage-specific transcriptional programs underlying migration of cortical neurons

**DOI:** 10.64898/2026.08.23.746411

**Authors:** Nitin Agnihotri, Ankita Jena, Manju Moorthy, Vijayalakshmi Bhat, Jonaki Sen

## Abstract

The laminar architecture of the mammalian neocortex depends on precise radial migration of newborn neurons to the appropriate cortical layers. This process is governed by the integration of extracellular signals with cell-intrinsic transcriptional programs. BMP signaling has been previously demonstrated to be essential for radial migration of late-born (E15.5) upper-layer cortical neurons. However, the gene expression programs downstream of BMP signaling that regulate this process remained unknown. To address this, we combined temporally targeted *in utero* electroporation with GeoMx Digital Spatial Profiling (DSP) to map BMP-responsive transcriptional programs in E15.5-born layer II/III neurons at two defined developmental timepoints: E17.5, when neurons actively migrate through the intermediate zone, and postnatal day 0 (P0), when they have completed migration and have attained their laminar position. BMP inhibition produced largely non-overlapping transcriptional changes at these two stages. At E17.5, chromatin-regulatory programs and ribosomal protein gene expression were collectively upregulated upon BMP inhibition. However, by P0, the same cohort of ribosomal genes exhibited downregulation while membrane lipid biosynthesis and synaptic specialization pathways became dominant, revealing a stage-dependent transcriptional switch. A subset of shared BMP-responsive genes was regulated in opposite directions at these two stages, which lent further support to the hypothesis that there is a temporal reorganization of BMP-dependent transcriptional outputs. We selected four candidates from among the BMP-responsive genes for functional studies, namely *Mfap4, Olfm2, Adora1,* and *Arpp21, which* belong to diverse functional categories, including extracellular matrix proteins, G protein-coupled receptors, secreted glycoproteins, and RNA-binding proteins. RNAi-mediated knockdown of all four candidates resulted in radial migration defects that closely phenocopied inhibition of BMP signaling, establishing these genes as functional effectors of the BMP signaling pathway regulating neuronal migration.

**Significance Statement:** The transcriptional programs through which BMP signaling regulates radial migration of upper-layer cortical neurons were completely unknown. By combining *in utero* electroporation with GeoMx spatial profiling at two developmental stages, this study maps BMP-responsive gene expression in defined migrating neurons *in vivo and* shows that BMP signaling engages largely non-overlapping transcriptional outputs that shift from chromatin and ribosomal programs during active migration to membrane lipid and synaptic programs at the laminar positioning stage. Further, four effector genes belonging to distinct molecular categories have been identified, each of which is essential for precise radial migration of E15.5-born neurons.

## Introduction

The laminar architecture of the mammalian cerebral cortex is assembled through precisely orchestrated developmental processes whereby neurons born at different time points migrate to distinct layers, acquire specific identities, and integrate into functional circuits. This gives rise to a highly ordered structure in which neurons born later occupy more superficial positions, a principle first demonstrated through birth-dating studies and later elaborated through the radial unit hypothesis, which proposes clonal columnar organization of cortical neurons (Angevine & Sidman, 1961; Rakic, 1974, 1988). Excitatory projection neurons in the cortex arise from radial glial progenitors lining the ventricular zone. These cells serve dual functions as neural stem cells as well as migratory scaffolds for new-born neurons that travel outward along radial glial processes to populate the cortical plate (Kriegstein & Alvarez-Buylla, 2009; Lui et al., 2011; Noctor et al., 2001). During this journey, new-born neurons pass through a multipolar phase in the intermediate zone before switching to a bipolar morphology to undergo glia-guided locomotion, a transition that requires precise control of cell polarity, cytoskeletal organization, and adhesive properties (Marín et al., 2010; Nadarajah et al., 2001; Noctor et al., 2004; Solecki, 2012; Tabata & Nakajima, 2003). Each of these steps depends on the integration of cell-intrinsic transcriptional programs with extracellular signaling inputs, which together ensure that neurons reach their correct laminar position (Ayala et al., 2007; Behar et al., 2001; Govek et al., 2011; Hippenmeyer et al., 2010; Marín & Rubenstein, 2001). When this coordination is disrupted, neurons stall or misposition, and cortical lamination is compromised, contributing to a spectrum of neurodevelopmental disorders such as lissencephaly, epilepsy, and autism spectrum disorders (Buchsbaum & Cappello, 2019; Guerrini & Dobyns, 2014).

Bone morphogenetic protein (BMP) signaling regulates multiple aspects of cortical development, including early dorsal telencephalic patterning, progenitor proliferation, and the switch between neurogenic and gliogenic competence (Furuta et al., 1997; Gomes et al., 2003; Gross et al., 1996; Panchision et al., 2001). BMP ligands and their signaling components are dynamically expressed in the developing cortex, where they regulate progenitor maintenance and the transition toward astroglial fates (Mabie et al., 1999; Miller & Gauthier, 2007; Saxena et al., 2018; Zhu et al., 1999). Previous work from our group demonstrated that BMP signaling specifically regulates the radial migration of embryonic day 15.5 (E15.5) born layer II/III cortical projection neurons (Saxena et al., 2018), extending its role beyond early patterning and fate specification. However, the downstream gene expression programs through which BMP signaling coordinates the changes in morphology, adhesion, and motility required for migration of this neuronal population remain uncharacterized. In addition, whether BMP signaling acts through a single transcriptional program or deploys distinct outputs at different time points along the migratory trajectory of upper-layer neurons also remained an open question.

Recent advances in single-cell and spatial transcriptomic approaches have substantially improved our understanding of cortical development, enabling high-resolution profiling of neuronal subtypes and their spatial organization within the developing brain. These studies have revealed extensive molecular heterogeneity among developing cortical neurons and have begun to define transcriptional trajectories associated with neuronal differentiation and circuit assembly (Di Bella et al., 2021; La Manno et al., 2021; Loo et al., 2019; Nowakowski et al., 2017). Despite this progress, a central challenge remains in linking extracellular signaling pathways to spatially and temporally resolved transcriptional programs within defined populations of migrating neurons. Majority of existing spatial transcriptomic studies have profiled unperturbed cortical tissue, which precludes the attribution of transcriptional changes to a defined signaling pathway, within a specific neuronal cohort along the migratory path.

To address this issue, one could envisage carrying out bulk transcriptomics of the embryonic mouse cortex after inhibition of a specific signaling pathway, such as BMP, by delivering dominant negative receptor expressing constructs through *in utero* electroporation. Since *in utero* electroporation can target only a small sub-population of cells within the cortex, this would dilute the effect of manipulation of BMP signaling, in the background of mostly unmanipulated cells. Alternatively, if one were to use Cre-lox-based strategies to knock out BMP ligands in the cortex, followed by bulk transcriptomics, this would avoid dilution but largely sacrifice spatiotemporal precision. Thus, none of the above-mentioned approaches could provide answers as to how a pathway such as BMP signaling modulates the gene expression landscape of a defined migrating neuronal population. Further, these approaches could not be used to determine if the transcriptional outputs of BMP signaling differ between the active migratory phase and the subsequent phase of laminar positioning.

Here, we have addressed this directly by combining *in utero* electroporation-based acute perturbation of the BMP pathway along with GeoMx Digital Spatial Profiling (DSP) in the developing mouse cortex. We map BMP-responsive gene expression programs in E15.5-born upper-layer cortical neurons, a neuronal population whose migration is dependent on active BMP signaling (Saxena et al., 2018), at two key developmental timepoints: embryonic day 17.5 (E17.5), when these neurons are actively migrating, and postnatal day 0 (P0), when migration is completing and laminar positioning is underway. Transcriptomic profiling of these populations revealed a pronounced stage-dependent shift in BMP-responsive transcriptional programs, with distinct pathway signatures at E17.5 and P0 and opposing regulation of certain gene cohorts across stages. Further, from among the differentially expressed genes identified through this approach, we shortlisted four candidate BMP-responsive genes-*Mfap4*, *Olfm2*, *Adora1*, and *Arpp21*, for functional characterization. These genes belonged to distinct molecular classes, yet functional perturbation of each of these genes reproduced the radial migration defects caused by inhibition of BMP signaling, implicating them as functional effectors rather than passive correlates. Taken together, our findings show that BMP-signaling shapes a temporally structured transcriptional program in upper-layer cortical neurons, one that shifts as these cells transition from active radial migration to laminar positioning. Moreover, this provides a spatially resolved, pathway-defined transcriptional framework for dissecting the molecular basis of upper-layer cortical neuron migration.

## Results

### 1) Spatial transcriptomic profiling captures stage-specific BMP-dependent transcriptional programs in migrating upper-layer cortical neurons

To map BMP-dependent transcriptional programs in E15.5-born layer II/III cortical neurons, we combined temporally targeted *in utero* electroporation (IUE) with GeoMx Digital Spatial Profiling (DSP) using the Mouse Whole Transcriptome Atlas (WTA). To inhibit BMP signaling cell-autonomously in upper-layer (layer II/III) progenitors, we electroporated a dominant-negative BMP receptor type Ib construct (pCAG-dnBMPR1b-IRES-GFP) in the developing cortex of timed-pregnant mice at embryonic day 15.5 (E15.5). The control construct, pCAG-IRES-GFP, was also electroporated in the mouse cortex at the same stage (Fig. 1A). The validation of dnBMPR1b-mediated downregulation of the BMP pathway and its effect on neuronal migration is described in the Materials and Methods and in Supplementary Fig. S1 (A, B). Electroporated cortices were harvested at two developmental timepoints: E17.5, when E15.5-born neurons are actively migrating through the intermediate zone, and postnatal day 0 (P0), when the majority of labeled neurons have completed radial migration and are undergoing laminar positioning within the cortical plate.

**Figure 1.**
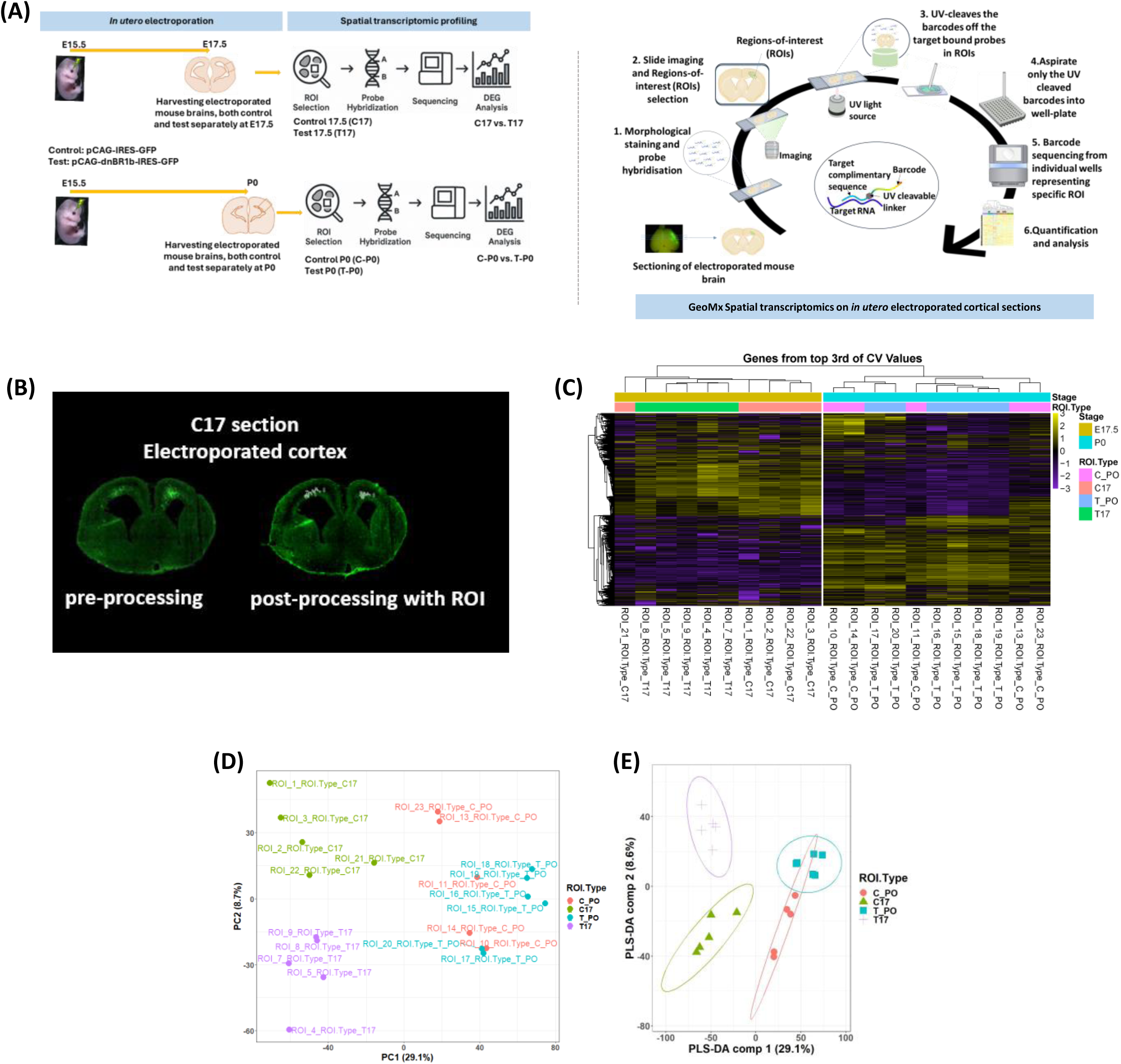
Spatial transcriptomic profiling captures stage-specific BMP dependent transcriptional programs in migrating upper-layer cortical neurons. **(A)** Experimental design. *In utero* electroporation (IUE) was performed at embryonic day 15.5 (E15.5) to introduce a dominant-negative BMP receptor type Ib construct (pCAG-dnBMPR1b-IRES-GFP; Test) or a control construct (pCAG-IRES-GFP) into cortical progenitors fated to generate upper-layer (layer II/III) neurons. Electroporated cortices were harvested at two developmental timepoints: E17.5, when E15.5-born neurons are actively migrating through the intermediate zone, and postnatal day 0 (P0), when radial migration is completing and neurons are undergoing laminar positioning within the cortical plate. Both control and dnBMPR1b-electroporated animals were collected at each timepoint, yielding four experimental groups: C17 (control, E17.5), T17 (dnBMPR1b, E17.5), C_PO (control, P0), and T_PO (dnBMPR1b, P0) and the schematic describing the workflow GeoMx Digital Spatial Profiling (DSP) using the Mouse Whole Transcriptome Atlas (WTA; 19,962-gene panel) performed on coronal cryosections from each group and the schematic representation of probe with gene specific barcodes attached to target complimentary region by UV cleavable linker. **(B)** Representative fluorescence images of coronal cortical sections from control-and dnBMPR1b-electroporated animals at E17.5, showing GFP signal (green) and DAPI counterstain. Manually delineated polygonal regions of interest (ROIs) are overlaid on GFP fluorescence; ROI boundaries were drawn based on GFP signal and cytoarchitectural landmarks to selectively capture the electroporated neuronal population within its spatial context. **(C)** Unsupervised hierarchical clustering heatmap of the top one-third most variable genes across all 23 Q3-normalised ROIs ranked by their coefficient of variation (CV). Columns represent individual ROIs, annotated by developmental stage (E17.5 or P0) and ROI type (C17, T17, C_PO, T_PO); rows represent genes. Colours represent row-scaled z-scores of Q3-normalised expression values. ROIs cluster by developmental stage as the dominant axis; within the E17.5 cohort, further separation by BMP inhibition status is visible. Within the P0 cohort, separation between groups is present but less pronounced, consistent with the more attenuated transcriptional response to BMP inhibition at this stage. **(D)** Principal component analysis (PCA) of Q3-normalised expression profiles across all 23 ROIs. PC1 (29.1% variance explained) segregates E17.5 from P0 ROIs; within the E17.5 cohort, PC2 (8.7% variance explained) shows partial separation between dnBMPR1b (T17) and control (C17) ROIs. Within the P0 cohort, groups are more closely positioned with partial overlap, consistent with the more attenuated transcriptional response to BMP inhibition at this stage. **(E)** PLS-DA (component 1 = 29.1%, component 2 = 8.6%) confirms distinct group separation across all four conditions and is shown for visualisation of group structure only; as a supervised method it was not used for statistical inference. Each point represents one ROI; colours indicate ROI type. IUE, in utero electroporation; DSP, Digital Spatial Profiling; WTA, Whole Transcriptome Atlas; PCA, principal component analysis; PLS-DA, partial least squares discriminant analysis; ROI, region of interest; Q3, upper-quartile normalisation. n = 5 ROIs per group (C17, T17, C_PO) and 6 ROIs (T_PO), Nuclei per ROI ranged from 66 to 575 across GFP-positive segments.

Coronal cryosections from control and dnBMPR1b-electroporated cortices at both stages were processed for GeoMx digital spatial profiling (Fig. 1A). Polygonal regions of interest (ROIs) were manually delineated based on GFP fluorescence and cytoarchitectural landmarks, enabling selective transcriptomic capture of electroporated populations within their native spatial and developmental context (Fig. 1B). In total, 23 ROIs were analyzed: five ROIs each from control E17.5 (C17), dnBMPR1b E17.5 (T17) and control P0 (C_P0) cortices, six ROIs from dnBMPR1b P0 (T_P0) cortices, and two GFP-negative ROIs one each E17.5 and P0 were included as internal negative references (Fig. 1B). ROIs were defined to encompass the GFP-positive zone; with nuclei count ranging from 66 to 575 per ROI, across all GFP-positive segments (Table 1). All ROIs met predefined area and signal thresholds and were retained for analysis. Segment-specific LOQ (limit of quantification) values ranged from 13.9 to 98.9 across all 23 ROIs; all segments met this threshold (Supplementary Fig. S2B). Following LOQ filtering, 9,550 genes detected above LOQ in at least 1% of segments were retained for downstream analysis. Gene detection rates ranged from 3.6% to 43.1% of the 19,962 genes in the panel; detection was lowest among C17 ROIs, none of which exceeded the 15% detection threshold, and uniformly highest among T_P0 ROIs, all six of which exceeded 29% (Supplementary Fig. S2 A, B). The higher detection rate in T_P0 ROIs was not attributable to greater nuclei counts since the T_P0 ROIs contained, on average, fewer nuclei than C17 ROIs (mean 144 vs 342 per ROI), yet substantially more genes were detected. For example, a T_P0 ROI with just 66 nuclei exceeded 36% gene detection, while the C17 ROI with the most nuclei (575) reached only 11.9% gene detection (Supplementary Fig. S2 A, B). This dissociation indicates that post-migratory neurons at P0, particularly those in which BMP signaling has been inhibited, engage a broader transcriptional program than neurons actively traversing the intermediate zone at E17.5, a difference that cannot be attributed to differences in ROI cellularity or sequencing depth. Unsupervised hierarchical clustering of the top one-third of the most highly variable genes confirmed that ROIs clustered coherently according to developmental stage and, within each stage, according to BMP inhibition status, demonstrating the reproducibility and internal consistency of the profiling across the replicates (Fig. 1C).

Following Q3 normalization, principal component analysis (PCA) revealed that PC1 (29.1% variance) segregated samples by developmental stage, with E17.5 ROIs clustering to the left and P0 ROIs to the right of the ordination space. Within the E17.5 cohort, dnBMPR1b and control ROIs displayed clear separation along PC2 (8.7% variance); while within the P0 cohort, the two groups were more closely positioned with partial overlap, indicating a more attenuated transcriptional response to BMP inhibition at this stage (Fig. 1D). Supervised partial least squares discriminant analysis (PLS-DA; component 1 = 29.1%, component 2 = 8.6%) confirmed distinct group separation across all four conditions; as a supervised method, this analysis was used for visualization rather than statistical inference (Fig. 1E). To define the cellular composition of the profiled ROIs at E17.5 and P0, we performed cell-type abundance deconvolution using SpatialDecon (Danaher et al., 2022) with custom gene expression profile matrices derived from stage-matched single-cell RNA-seq datasets (GSE153162 for E17.5 and GSE123335 for P0). The reference profiles were generated from the corresponding published single-cell transcriptomic studies. At E17.5, migrating neurons represented the predominant cell-type signature across GFP-positive ROIs in both control and dnBMPR1b conditions, with mean abundance scores of 103 and 111, respectively, confirming enrichment of the intended neuronal population (Table 2; Supplementary Fig. S2D). At P0, the profiled ROIs showed prominent Layer II–IV cortical neuron signatures, with mean abundance scores of 60.4 and 36.8 in control and dnBMPR1b conditions, respectively, consistent with the upper-layer neuronal population analyzed at this stage (Table 3; Supplementary Fig. S2D). The broader cell-type abundance profiles for the analyzed ROIs are shown in Supplementary Fig. S2D.

Collectively, these data demonstrated that GeoMx spatial profiling of IUE-targeted cortices captures spatially resolved transcriptional variation associated with BMP pathway perturbation in E15.5-born upper-layer neurons at two developmental stages. This stage-dependent transcriptional divergence, more pronounced at E17.5 and distinct in character at P0, motivates stage-resolved interrogation of BMP-regulated molecular programs, which we characterized as described in the following sections.

### 2) BMP signaling regulates stage-specific, largely non-overlapping transcriptional programs in E15.5 born migrating upper-layer neurons

To distinguish transcriptional changes during active migration from those associated with post-migratory laminar positioning, we performed differential expression (DE) analysis comparing dnBMPR1b and control GFP-positive ROIs separately at E17.5 and P0 (Fig. 2A).

**Figure 2.**
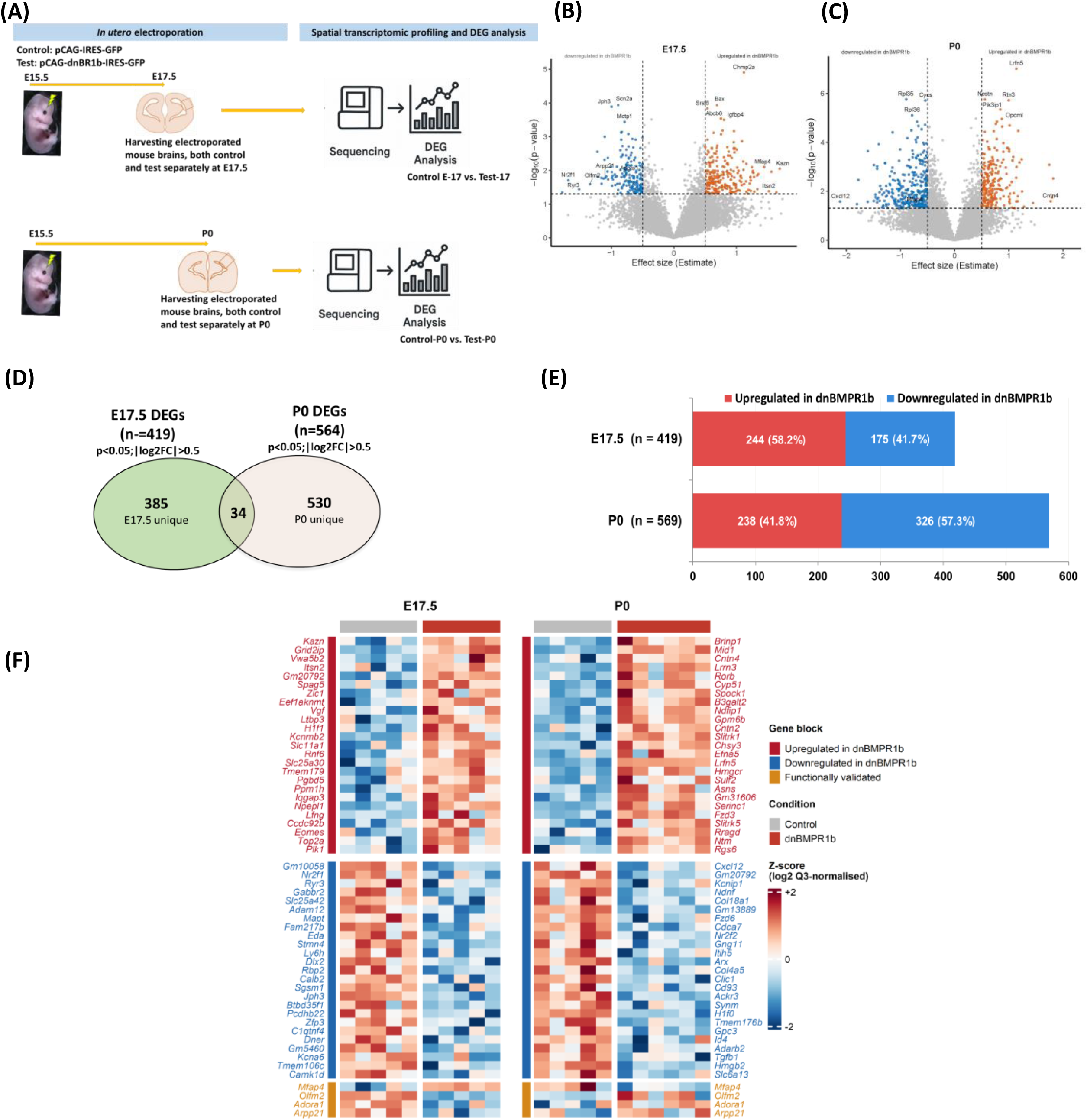
BMP signaling regulates stage-specific, largely non-overlapping transcriptional programs in E15.5 born migrating upper-layer neurons. **(A)** Schematic of the differential expression (DE) analysis strategy of E17.5 and P0 cortices that were electroporated with test or control plasmids at E15.5. Linear mixed model (LMM) analysis was performed separately for the E17.5 comparison (T17 vs C17) and P0 comparison (T_PO vs C_PO) using the GeoMxTools pipeline. Genes meeting p < 0.05 and |log₂FC| > 0.5 were retained as the fold-change-filtered gene set for all downstream analyses. Y-chromosome-linked transcripts (*Ddx3y*, *Kdm5d*, *Eif2s3y*, *Uty*, *Gm21783*) were excluded prior to all downstream analyses. **(B)** Volcano plot of differential expression at E17.5 (T17 vs C17). x-axis, effect size estimate; y-axis, - log₁₀(p-value). Horizontal dashed line, p = 0.05; vertical dashed lines, |log₂FC| = 0.5. Points coloured by direction and significance: red, upregulated (log₂FC > 0.5, p < 0.05); blue, downregulated (log₂FC < -0.5, p < 0.05); green, nominally significant but below fold-change threshold; grey, not significant. Of 644 genes with p < 0.05, 424 met the fold-change threshold (249 upregulated, 175 downregulated). Five Y-chromosome-linked transcripts (*Ddx3y*, *Kdm5d*, *Eif2s3y*, *Uty*, *Gm21783*), all upregulated, were excluded from downstream analyses, yielding 639 genes at p < 0.05 and 419 at the fold-change threshold (244 upregulated, 175 downregulated). Full E17.5 DEG lists are provided in supplementary table 4. **(C)** Volcano plot of differential expression at P0 (T_PO vs C_PO). Axis and colour conventions as in (B). Of **1,116** genes with p < 0.05, 569 met the fold-change threshold (238 upregulated, 331 downregulated). Five Y-chromosome-linked transcripts (*Ddx3y*, *Kdm5d*, *Eif2s3y*, *Uty*, *Gm21783*), all downregulated at P0, were excluded from all downstream analyses, yielding 1,111 genes at p < 0.05 and 564 at the fold-change threshold (238 upregulated, 326 downregulated), with a bias toward downregulation. Full P0 DEG lists are provided in supplementary table 5. **(D)** Venn diagram showing the overlap between fold-change filtered DEG sets at E17.5 (419 genes) and P0 (564 genes). Of 949 genes in the combined non-redundant set, 34 were shared between stages (3.6%). Of these 34 genes, 13 were regulated concordantly at both timepoints (4 upregulated at both stages, 9 downregulated at both stages) and 21 showed opposing regulation, of which 17 were upregulated at E17.5 and downregulated at P0. This directional bias (81% of reversals following the same Up → Down pattern) is consistent with a genuine transcriptional switch rather than stochastic stage-to-stage variation. The shared genes with directional annotations are listed in supplementary table 8. **(E)** A horizontal stacked bar chart depicting proportion of upregulated and downregulated genes. Post Y-chromosome-exclusion, at E17.5: 419 genes total (244 upregulated, 58.2%; 175 downregulated, 41.7%). P0: 564 genes total (238 upregulated, 41.6%; 326 downregulated, 57.3%) **(F)** Heatmap of top fold-change-filtered DEGs at E17.5 (left) and P0 (right) across GFP-positive ROIs. At each stage, the 25 most upregulated and 25 most downregulated genes by effect size are shown (Y-chromosome-linked genes excluded). Four functionally validated candidates (*Mfap4*, *Olfm2*, *Adora1*, *Arpp21*) are displayed as a separate block (amber annotation bar). Genes are ordered by effect size within each block; columns represent individual ROIs (E17.5: n = 5 C17, 5 T17; P0: n = 5 C_PO, 6 T_PO) grouped by condition. Colours represent row-scaled z-scores of log₂ Q3-normalised expression values. Gene labels: red, upregulated in dnBMPR1b; blue, downregulated in dnBMPR1b; amber, functionally validated candidates. Full DEG heatmaps are provided in supplementary file. DEG, differentially expressed gene; LMM, linear mixed model; FC, fold change. Significance threshold: p < 0.05, |log₂FC| > 0.5. n = 5 ROIs per group (C17, T17, C_PO) and 6 ROIs (T_PO).

BMP inhibition produced stage-dependent transcriptional effects, with a more acute transcriptional response at E17.5 and broader gene-level differential expression at P0. At E17.5, 644 genes were significantly altered (p < 0.05), of which 424 met the fold-change threshold (|log₂FC| > 0.5), comprising 249 upregulated and 175 downregulated genes (Fig. 2B, Table 4). Five Y-chromosome-linked transcripts (Ddx3y, Kdm5d, Eif2s3y, Uty, Gm21783) present among the upregulated genes were excluded from all downstream analyses, yielding 639 genes (p<0.05) and 419 FC-filtered genes (244 upregulated, 175 downregulated). On the other hand, at P0, 1,116 genes were significantly differentially expressed (p < 0.05), of which 569 met the fold-change threshold (238 upregulated, 331 downregulated), showing a bias toward downregulation (Fig. 2C, Table 5). After excluding five Y-chromosome-linked transcripts present among the downregulated genes from downstream analyses yielding 1,111 genes (p < 0.05) and 564 FC-filtered genes (238 upregulated, 326 downregulated) (Fig. 2E). The greater number of differentially expressed genes at P0 likely reflects broader transcriptional engagement during post-migratory maturation rather than a stronger perturbation effect per se, consistent with the more attenuated BMP-dependent separation observed in PCA at this stage (Fig. 1D). Individual gene-level FDR values among fold-change-filtered DEGs range from 0.001 to 0.43 at P0 and from 0.12 to 0.74 at E17.5, the latter reflecting the statistical regime inherent to small-cohort spatial profiling (n = 5-6 ROIs per group) and the absence of multiple-testing correction. Individual genes mentioned throughout should be treated as candidate DEGs for functional follow-up rather than as FDR-controlled targets.

Among the genes upregulated upon BMP inhibition at E17.5 were *Kazn*, encoding kazrin (estimate=+1.69, p=0.009), a periplakin-interacting scaffolding protein that regulates cytoskeletal organization and cell adhesion in epithelial cells (Sevilla et al., 2008) and promotes dynein/dynactin-dependent endosomal trafficking (Hernandez-Perez et al., 2023), and *Itsn2*, encoding intersectin-2 (estimate=+1.52, p=0.042), an evolutionarily conserved multidomain scaffold involved in endocytosis and actin-associated membrane trafficking (Tsyba et al., 2011). Downregulated genes at this stage included *Nr2f1* (also known as COUP-TFI) (estimate=-1.69, p=0.020), a nuclear receptor that regulates cortical neuronal migration and laminar specification (Alfano et al., 2011), and *Ryr3* (estimate=-1.52, p=0.036), encoding a ryanodine receptor calcium channel expressed in murine brain (Giannini et al., 1995). The downregulation of *Nr2f1* is consistent with a role for COUP-TFI in promoting radial migration of callosal projection neurons by repressing *Rnd2* (Alfano et al., 2011), a Rho GTPase required for the multipolar-to-bipolar transition during migration (Heng et al., 2008). The reduced expression of *Nr2f1* following BMP inhibition suggests that BMP signaling acts through transcriptional regulators such as NR2F1 to coordinate the cytoskeletal dynamics underlying radial migration of upper-layer neurons. Together, these changes are consistent with disruption of cytoskeletal organization, membrane trafficking, and transcriptional control mechanisms that are central to the polarity transition and directed radial locomotion of migrating neurons.

Among nominally significant individual genes at P0, *Cntn4* (contactin-4 *or Big2*), an immunoglobulin superfamily adhesion molecule implicated in axon guidance and cortical circuit formation (Kaneko-Goto et al., 2008; Osterhout et al., 2015), was upregulated (Estimate = +1.77, p = 0.026; FDR = 0.33), and *Cxcl12* (SDF-1) was downregulated (Estimate = -2.12, p = 0.027; FDR = 0.33). However, both these genes represent candidate findings at this sample size and require independent validation. Taken together, these observations are consistent with BMP signaling at P0 influencing intercellular adhesion and the emerging synaptic environment as neurons complete migration and consolidate laminar position.

Direct comparison of the fold-change filtered DEG sets at E17.5 and P0 revealed minimal overlap, with only 34 genes shared between the two stages (3.6% of the combined non-redundant set; Fig. 2D). Of these, 13 were regulated in the same direction at both timepoints (4 upregulated and 9 downregulated), while the remaining 21 showed opposing regulation (Fig. 2E). The directional bias within this overlap was pronounced: 17 of these 21 genes were upregulated at E17.5 and downregulated at P0. Although the absolute number of shared genes is small, the consistency of this pattern-the same directional logic (upregulated at E17.5, downregulated at P0) governing 17 of 21 reversals suggests that the limited overlap reflects a genuine transcriptional switch rather than stochastic stage-to-stage variation. Unsupervised clustering of the top differentially expressed genes at each stage segregated samples by BMP inhibition status, with dnBMPR1b and control ROIs forming distinct clusters at both E17.5 and P0, further confirming the robustness and stage-specificity of the identified transcriptional signatures (Fig. 2F). Among the highest-ranked genes driving E17.5 upregulation were *Kazn* and *Itsn2*, consistent with the cytoskeletal and membrane trafficking programs identified by GSEA. *Mfap4* ranked fifth among upregulated genes at E17.5 yet appeared in the downregulated block at P0, illustrating at the single-gene level the directional reversal that characterizes the overlapping DEG set at the two stages (Fig. 2F). Together, these findings indicate that BMP-responsive transcriptional programs are largely non-overlapping across developmental stages and, where overlap exists, are regulated in opposing directions in 81% of cases (17 of 21 shared genes with opposing regulation), consistent with BMP signaling contributing to the temporal coordination of gene expression as neurons transition from active radial migration to post-migratory maturation. BMP signaling, therefore, governs largely distinct, stage-appropriate transcriptional programs as upper-layer neurons progress from the intermediate zone at E17.5 to laminar positioning at P0.

### 3) BMP-regulated transcriptional programs undergo a stage-dependent switch from chromatin-ribosomal to lipid biosynthesis and synaptic pathways

To identify the biological processes associated with BMP-dependent transcriptional changes at each stage, we performed GSEA separately on the entire ranked lists from E17.5 and P0. At E17.5, BMP inhibition was associated with significant enrichment across 200 gene sets (FDR < 0.05; Fig. 3A; Table 6, spanning GO biological process, cellular component, and molecular function categories, as well as reactome pathways. Within the GO biological process category, 88 gene sets reached significance (80 positively enriched, 8 negatively enriched). The most strongly positively enriched categories included chromosome condensation (NES = 2.08), mitotic sister chromatid segregation (NES = 2.04), cytoplasmic translation (NES = 2.00), chromosome organization (NES = 1.94), and mRNA processing (NES = 1.54). These signatures were accompanied by elevated expression of genes, including *Plk1*, *Ube2c*, *Bub1b*, and *Spag5* in dnBMPR1b ROIs relative to controls. Although these genes are canonically associated with cell division, emerging evidence indicates that they also function in the post-mitotic neurons; for example, Aurora kinase B has been shown to regulate axonal outgrowth independently of its mitotic role (Gwee et al., 2018), and APC/C-CDH1 components regulate the proteasomal clearance of chromatin-associated mitotic factors during neuronal differentiation (Ledvin et al., 2023). The enrichment of these gene products in GFP-positive ROIs therefore reflects BMP-dependent regulation of non-canonical chromatin or cytoskeletal programs in migrating neurons, though a contribution from residual progenitor populations cannot be entirely excluded (Table 6).

**Figure 3.**
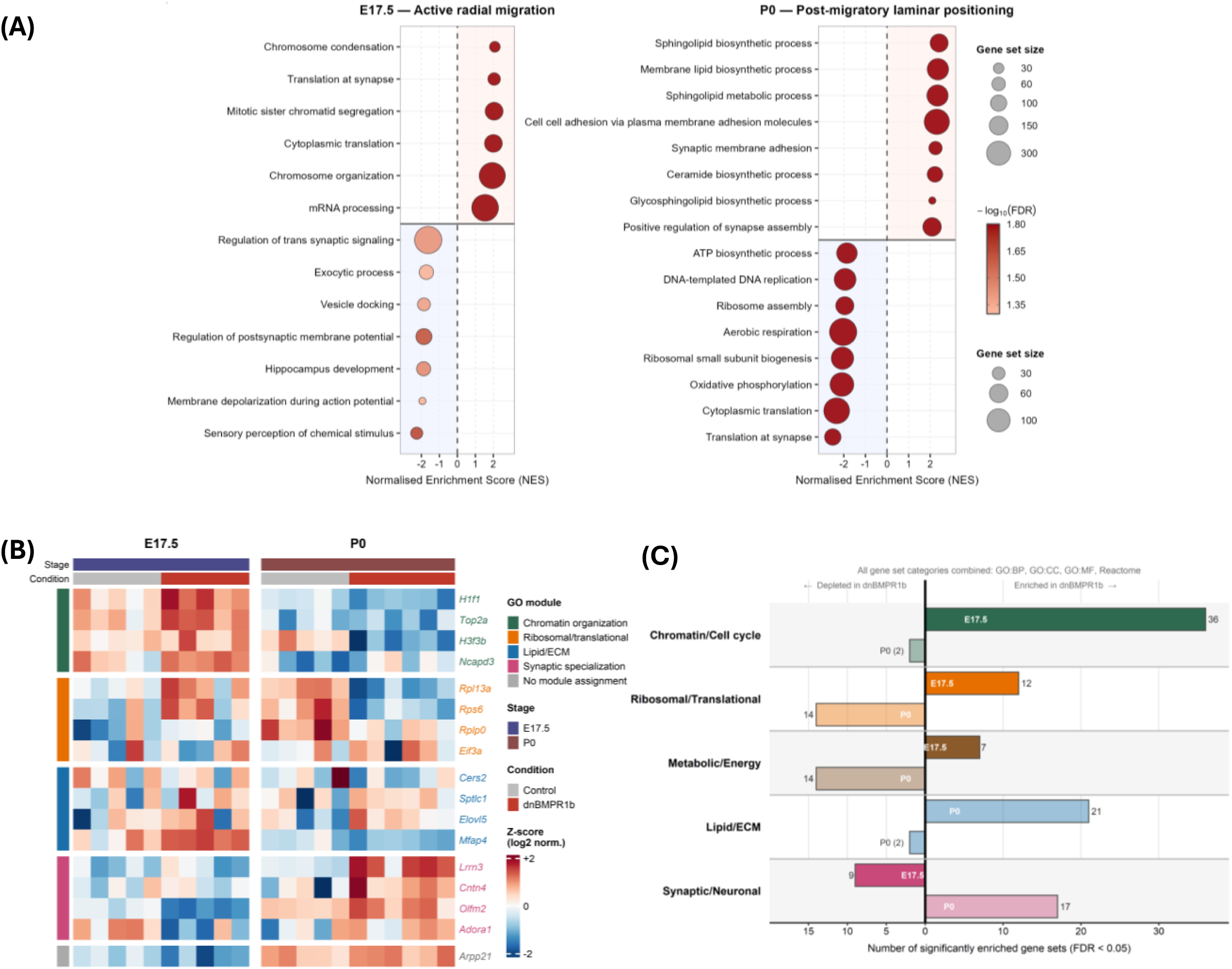
BMP-regulated transcriptional programs undergo a stage-dependent switch from chromatin-ribosomal to lipid biosynthesis and synaptic pathways. (A) Bubble plots showing selected significantly enriched GO Biological Process gene sets at E17.5 (left) and P0 (right), from gene set enrichment analysis (GSEA) at each stage. Each panel is split horizontally: the upper section shows gene sets positively enriched in dnBMPR1b ROIs relative to controls (NES > 0); the lower section shows negatively enriched gene sets (NES < 0). Position on the x-axis indicates the normalized enrichment score (NES); bubble size represents gene set size; bubble colour represents significance (-log₁₀(FDR)), with darker red indicating greater significance. The dashed vertical line marks NES = 0. Gene sets shown represent a curated selection of the top enriched categories by absolute NES from the full set of 200 significantly enriched gene sets at E17.5 (GO: BP = 88, GO:CC = 42, GO:MF = 13, Reactome = 57) and 321 at P0 (GO: BP = 161, GO:CC = 78, GO:MF = 22, Reactome = 60). Complete GSEA results are provided in Tables S6 (E17.5) and S7 (P0). (B) Non-candidate representative genes were selected as leading-edge members of FDR-significant gene sets within each module category (GO: BP, FDR < 0.05); ceramide and sphingolipid biosynthetic enzymes (Cers2, Sptlc1, Elovl5) and Eif3a were additionally included as biologically representative members of their respective enriched modules in the represented heatmap. Three validated candidates (Mfap4, Olfm2, Adora1) are placed within their biologically cognate modules, one candidate (Arpp21) shown without module assignment, Genes are not clustered; order within each module is fixed by effect size rank within the leading edge. (C) Diverging bar chart showing the number of significantly enriched GO Biological Process, GO Cellular Component, GO Molecular Function, and Reactome gene sets (FDR < 0.05) at E17.5 and P0, grouped by broad functional module. Bars extending to the right (positive direction) indicate gene sets enriched in dnBMPR1b ROIs; bars extending to the left (negative direction) indicate gene sets depleted in dnBMPR1b ROIs. Dark bars, E17.5; light bars, P0. At E17.5, BMP inhibition is predominantly associated with enrichment of chromatin/cell cycle programs (36 gene sets positively enriched), ribosomal/translational programs (12 positively enriched), and metabolic/energy programs (7 positively enriched), with reciprocal depletion of synaptic/neuronal programs (9 negatively enriched). At P0, the pattern shifts markedly: lipid/ECM programs become the dominant positively enriched module (21 gene sets), synaptic/neuronal programs switch from depleted to enriched (17 gene sets), and ribosomal/translational programs reverse to become predominantly depleted (14 gene sets negatively enriched), consistent with the stage-dependent transcriptional shift.

In addition, reactome pathway analysis revealed positive enrichment of epigenetic regulatory programs, including epigenetic regulation of gene expression (NES = 2.07), PRC2-mediated histone methylation (NES = 2.08), and formation of senescence-associated heterochromatin foci (NES = 2.09). These reactome pathways indicate heterochromatin assembly programs rather than neuronal senescence, which is consistent with active transcriptional remodeling during the migratory phase. This indicated that BMP inhibition during active radial migration is accompanied by upregulation of metabolic programs, which is consistent with the elevated translational capacity, apparent from the ribosomal protein gene enrichment at this stage. Cytoplasmic translation (NES = +2.00, FDR = 0.016) and the translation at synapse gene sets (NES = +2.04, FDR = 0.016) were also positively enriched at E17.5 in the ROIs where dnBMPR1b was overexpressed. Notably, the ‘translation at synapse’ term here reflects the broad assignment of ribosomal structural genes to synaptic translation processes in GO annotation, which may not be evidence of preferential synaptic translation. The leading-edge genes for both sets are composed predominantly of structural ribosomal protein genes of the Rpl and Rps families; all 25 leading-edge genes for the translation at synapse set encode ribosomal proteins or ribosomal fusion proteins (including Uba52/RPL40) with no canonical synaptic translation regulators present in either leading edge. These enrichments reflect a coordinated increase in ribosomal protein gene expression in dnBMPR1b ROIs at E17.5, consistent with an elevated ribosomal/translational gene-expression program during the migratory phase (Fig. 3A, Table 6).

In contrast, gene sets associated with mature neuronal physiology were negatively enriched at E17.5 (FDR < 0.05), including sensory perception of chemical stimulus (NES = -2.26), membrane depolarization during action potential (NES = -1.94), hippocampus development (NES = -1.87), regulation of postsynaptic membrane potential (NES = -1.86), vesicle docking (NES = -1.86), exocytic process (NES = -1.72), and regulation of trans-synaptic signaling (NES = -1.62) (Fig. 3A). This pattern of negative enrichment is consistent with the transcriptionally and functionally immature state of neurons traversing the intermediate zone, which have not yet established mature electrophysiological properties or synaptic connectivity.

Interestingly, at P0, the transcriptional landscape shifted markedly, with 321 gene sets significantly enriched (FDR < 0.05) across all pathway categories compared with 200 at E17.5, consistent with the larger number of differentially expressed genes at this stage and the broader maturation programs engaged as neurons reach the cortical plate (Fig. 3A; Table 7). Within the GO biological process category, 161 gene sets reached significance. BMP-responsive genes were strongly enriched for pathways linked to membrane lipid biosynthesis and synaptic specialization (Fig. 3A; Table 7, with the most highly enriched categories including sphingolipid biosynthetic process (NES = 2.41), membrane lipid biosynthetic process (NES = 2.36), sphingolipid metabolic process (NES = 2.34), cell-cell adhesion via plasma membrane adhesion molecules (NES = 2.30), synaptic membrane adhesion (NES = 2.24), ceramide biosynthetic process (NES = 2.22), glycosphingolipid biosynthetic process (NES = 2.10), and positive regulation of synapse assembly (NES = 2.09). Consistent with these categories, Lrrn3 (Nlrr3), a brain-enriched LRR transmembrane protein implicated in neural circuit development (Kochunov et al., 2013; Taniguchi et al., 1996), met the applied threshold (p < 0.05, |log₂FC| > 0.5) and was a leading-edge gene in nine FDR-significant synapse assembly and junction organization gene sets at P0.

Reactome analysis further identified enrichment for Eph-ephrin-mediated cell repulsion (NES = 1.96, FDR = 0.018), a pathway directly relevant to cortical neuron positioning and dendritic targeting. Consistent with these biological process enrichments, GO cellular component analysis at P0 identified strong positive enrichment of synaptic membrane compartments, including presynaptic membrane (NES = 2.22) and postsynaptic membrane (NES = 2.23), providing orthogonal spatial support for BMP-dependent membrane specialization at this stage (Fig. 3A; Table 7). These enrichments are consistent with a possible role for BMP signaling in supporting membrane remodeling and early circuit assembly as neurons complete migration and begin to mature within the cortical plate.

In parallel, a broad set of metabolic and translational programs was negatively enriched at P0 following BMP inhibition (FDR < 0.05). The most strongly depleted categories included translation at synapse (NES = -2.51), cytoplasmic translation (NES = -2.33), oxidative phosphorylation (NES = - 2.09), ribosomal small subunit biogenesis (NES = -2.06), aerobic respiration (NES = -2.03), DNA-templated DNA replication (NES = -1.94), ribosome assembly (NES = -1.95), and ATP biosynthetic process (NES = -1.86). The same ribosomal protein gene cohort that was collectively upregulated at E17.5 drives the negative enrichment of translational gene sets at P0: 22 of the 25 E17.5 leading-edge genes for the translation at synapse set are present in the P0 leading edge (88% overlap), and 39 of 57 cytoplasmic translation leading-edge genes are shared (Fig. 3A; Table 7). This stage-dependent reversal, with ribosomal protein genes collectively enriched in dnBMPR1b ROIs at E17.5 but depleted at P0, indicates that BMP inhibition has opposing effects on ribosomal gene-expression programs across developmental stages as neurons transition from active migration to post-migratory maturation within the cortical plate.

A heatmap of representative genes within these enriched categories revealed coordinated, stage-dependent regulation across GFP-positive ROIs (Fig. 3B). Grouping significantly enriched gene sets (FDR < 0.05) into broad functional modules confirmed that at E17.5, BMP inhibition is predominantly associated with chromatin/cell cycle and ribosomal/translational programs, whereas at P0 lipid/ECM and synaptic/neuronal modules became the dominant positively enriched categories, with ribosomal/translational programs reversing to net depletion, consistent with the stage-dependent transcriptional shift described above (Fig. 3C).

Thus, BMP signaling appears to be associated with distinct, largely non-overlapping transcriptional programs that differ fundamentally across developmental contexts. At E17.5, during active radial migration, BMP activity correlates with chromatin-associated and epigenetic programs, elevated ribosomal protein gene expression, and metabolic/energy programs, while electrophysiological and synaptic signaling programs associated with mature neurons are negatively enriched. By P0, BMP-associated transcriptional programs shift toward membrane lipid biosynthesis, cell adhesion, and synaptic specialization, consistent with the transition to post-migratory maturation within the cortical plate.

### 4) BMP-responsive candidate genes are expressed in the developing cortex and show stage-dependent regulation consistent with BMP–SMAD transcriptional control

Building on the stage-resolved transcriptomic analysis described above, we sought to identify candidate downstream effectors of BMP signaling for functional validation. Candidates were selected from the E17.5 DEG set and prioritized according to the following criteria: 1) meeting the applied threshold (p < 0.05, |log₂FC| > 0.5); 2) exhibiting detectable cortical expression confirmed by RNA *in situ* hybridization; 3) representing diverse functional classes; and, where available, 4) prior evidence linking the gene or its molecular class to migration, polarity, dendritogenesis, or neurodevelopmental disease. We deliberately extended the search beyond canonical migration regulators to include extracellular matrix proteins, G-protein coupled receptors, secreted glycoproteins, and RNA-binding proteins.

We performed RNA *in situ* hybridization for a panel of DEGs in wild-type mouse cortices at E15.5 (Fig. S3). Several genes from the candidate list did not yield detectable signal, likely reflecting false-positive DEG calls, the sensitivity limitations of chromogenic *in situ* hybridization relative to spatial transcriptomic profiling, or both, and were not pursued further. Among genes yielding robust and reproducible cortical expression, *Mfap4,* a significantly upregulated DEG at E17.5, showed a strong signal in the ventricular zone (VZ) and subventricular zone (SVZ), with lower expression in the intermediate zone (IZ) following a medial-to-lateral gradient. Although the expression extended into the lower intermediate zone, expression in the cortical plate (CP) expression was minimal (Fig. 4A). From the set of downregulated genes, *Olfm2* was found to be expressed robustly in the VZ and SVZ with lower signal in the CP (Fig. 4B), *Adora1* transcripts were observed to be concentrated in the CP, with lower levels across the VZ, SVZ, and IZ (Fig. 4C) and *Arpp21* exhibited a similar CP-predominant distribution with low level of expression in lower cortical regions (Fig. 4D). These patterns place all four genes within the developing cortex at a stage and region when upper-layer neurons are actively migrating, consistent with their potential roles in this process. Other DEGs whose expression was also detected in the developing cortex are listed in Fig. S3.

**Figure 4:**
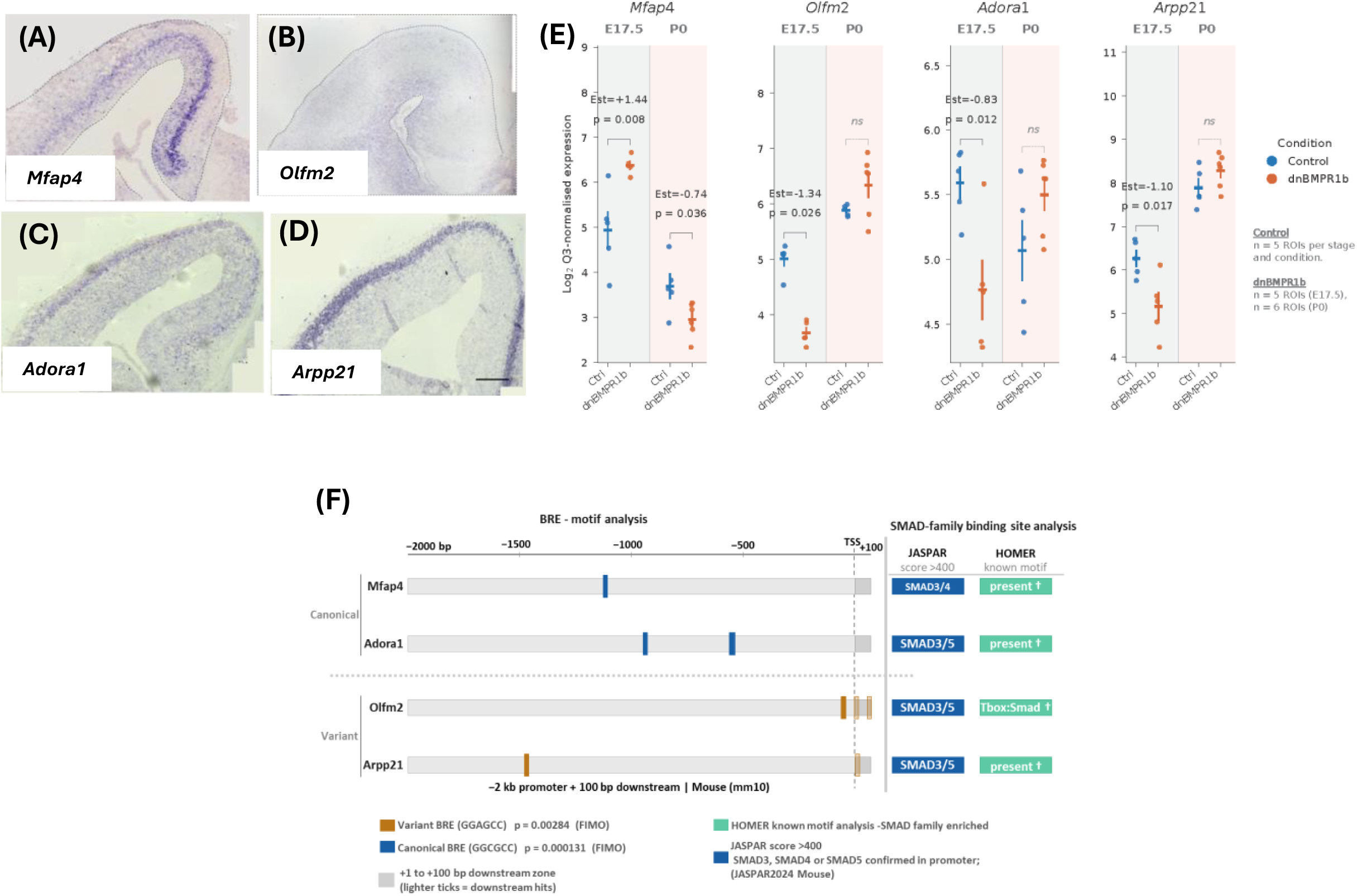
BMP-responsive candidate genes are expressed in the developing cortex and show stage-dependent regulation consistent with BMP–SMAD transcriptional control. **(A–D)** RNA in situ hybridization using digoxigenin-labelled antisense riboprobes in wild-type E15.5 coronal cortical sections, corresponding to the time of *in utero* electroporation. **(A)** *Mfap4*: strong signal in the VZ and SVZ, with lower expression in the IZ and a medial-to-lateral gradient; minimal CP expression. **(B)** *Olfm2*: robust signal in the VZ and SVZ, with lower expression in the CP. **(C)** *Adora1*: concentrated in the CP, with lower levels across VZ, SVZ, and IZ. **(D)** *Arpp21*: CP-predominant distribution. Scale bar:100 µm. **(E)** Q3-normalised GeoMx expression levels of four BMP-responsive candidates-*Mfap4*, *Olfm2*, *Adora1*, and *Arpp21* - across GFP-positive ROIs from control (Ctrl) and dnBMPR1b-electroporated cortices at E17.5 and P0. Each point represents one ROI; bars show mean ± SEM. Effect size estimates and p-values from LMM analysis are indicated. *Mfap4* was upregulated in dnBMPR1b ROIs at E17.5 (Estimate = +1.44, p = 0.008) and downregulated at P0 (Estimate = -0.74, p = 0.036), exhibiting a stage-dependent reversal. *Olfm2* (Estimate = -1.34, p = 0.026), *Adora1* (Estimate = -0.83, p = 0.012), and *Arpp21* (Estimate = -1.10, p = 0.017) were each downregulated in dnBMPR1b ROIs at E17.5; none showed statistically significant differential expression at P0 under the applied threshold (p < 0.05, |log₂FC| > 0.5). LMM, linear mixed model. **(F)** Positional map of BMP-responsive element (BRE) motifs and SMAD-family transcription factor binding site predictions within the 2-kb promoter region and 100 bp downstream of *Mfap4*, *Olfm2*, *Adora1*, and *Arpp21*. Left panel: horizontal bars represent the genomic window screened for each gene (-2000 to +1 bp relative to the transcription start site [TSS], grey; +1 to +100 bp downstream, dark grey). Vertical ticks/bars indicate BRE motif hit positions identified by FIMO (MEME Suite v5.5; meme-suite.org) using the canonical GC-rich BRE consensus (GGCGCC; blue ticks; p = 0.000131) and a BRE variant sequence (GGAGCC; amber ticks; p = 0.00284) as query motifs against the submitted promoter and downstream sequences. Tick positions are mapped proportionally from FIMO coordinate output; lighter amber ticks denote hits within the downstream (+1 to +100 bp) window. Genes are grouped by BRE motif type: canonical BRE (GGCGCC) identified in *Mfap4* (1 site) and *Adora1* (3 sites); variant BRE (GGAGCC) identified in *Olfm2* (1 promoter site, 2 downstream sites) and *Arpp21* (1 promoter site, 1 downstream site). Right panel: summary of SMAD-family transcription factor binding site predictions. JASPAR column: presence of SMAD-family motifs (SMAD3, SMAD4, or SMAD5) at JASPAR score >400, identified using the JASPAR TFBS extraction tool (jaspar.elixir.no) against the JASPAR2024 Mouse database within the same genomic windows. HOMER column: SMAD-family motif enrichment identified by known motif analysis (HOMER findMotifs.pl; Mouse genomic background). For *Olfm2*, the composite Tbox: Smad motif was the top-ranked known motif. HOMER analysis was performed on a single target sequence per gene; enrichment statistics are therefore non-discriminating and are presented as qualitative support only. No SMAD-family motifs at score >400 were identified in any downstream (+1 to +100 bp) region across all four genes. BRE, BMP-responsive element; TSS, transcription start site; FIMO, Find Individual Motif Occurrences; HOMER, Hypergeometric Optimization of Motif EnRichment.

We next examined spatial expression of all four candidates within the GeoMx dataset by visualizing Q3-normalized signal intensities across GFP-positive ROIs from control and dnBMPR1b-electroporated cortices at E17.5 and P0 (Fig. 4E). Consistent with the DEG analysis, Mfap4 expression was higher in dnBMPR1b ROIs at E17.5 (Estimate = +1.44, p = 0.008) and lower at P0 (Estimate = -0.74, p = 0.036), exhibiting a reversal in direction between stages. On the other hand, the expression of Olfm2, Adora1, and Arpp21 was reduced in dnBMPR1b ROIs at E17.5 (Estimates: - 1.34, -0.83, -1.10; p = 0.026, 0.012, 0.017 respectively), indicating BMP signaling normally supports their expression during active migration (Fig. 4E). With the exception of *Mfap4,* none of the other three genes exhibited statistically significant differential expression at P0 under the applied threshold. Interestingly, expression trends in dnBMPR1b ROIs at P0 for all three genes showed sub-threshold directional reversals, consistent with the stage-dependent pattern described in section 2.

To assess whether the identified candidates carry promoter features consistent with BMP-SMAD transcriptional control, we screened the 2-kb promoter regions and 100 bp downstream sequences of all four genes for BMP-responsive elements (BREs) (Korchynskyi & ten Dijke, 2002). Canonical GC-rich BREs (GGCGCC) were identified within the screened promoter regions of Mfap4 and Adora1 (p < 0.0005), while a variant BRE motif (GGAGCC) was identified within the screened promoter and downstream regions of Olfm2 and Arpp21 (p < 0.005) (Fig. 4F, Table 9). Computational screening of predicted transcription factor binding sites using JASPAR2024 database identified SMAD-family motifs (SMAD3, SMAD4, and/or SMAD5; score >400) in the screened regions of all four genes. Consistent with this, HOMER known motif analysis led to identification of SMAD-family motifs in all four screened regions, with the composite Tbox: Smad motif ranking as the top-ranked known motif for Olfm2. Further, a SMAD2-binding motif (MA1964.2) was additionally identified in the screened region of Mfap4 by *de novo* motif analysis. Together, these findings indicate that screened promoter and downstream regions of all four candidate genes contain features consistent with potential SMAD-mediated transcriptional regulation downstream of BMP signalling (Fig.4F, Table 10). Based on confirmed cortical expression, statistically supported differential regulation, stage-specific transcriptional context, and coverage of four distinct molecular classes, we selected *Mfap4, Olfm2, Adora1, and Arpp21* for RNAi-mediated functional studies to assess if they are involved in regulating migration of E15.5 born neurons.

### 5) RNAi-mediated knockdown of BMP-responsive candidate genes disrupts radial migration of upper-layer cortical neurons

To determine whether the BMP-responsive candidate genes identified through spatial transcriptomics are indeed necessary for radial migration, we employed a loss-of-function strategy using a microRNA-based RNAi construct delivered via *in utero* electroporation (Hand & Polleux, 2011; Smith et al., 2009). Gene-specific miRNA sequences targeting the 3′ UTR of each candidate were designed and cloned into the pRmiR vector (Smith et al., 2009), which co-expresses GFP to mark transfected cells (Fig 5A). Constructs were *in utero* electroporated into the developing cortices of timed-pregnant mice at E15.5, targeting progenitors fated to generate upper-layer (layer II/III) cortical neurons. The empty pRmiR vector served as the control in all experiments (Udaykumar et al., 2023). Electroporated cortices were analyzed at two developmental timepoints: E17.5, when E15.5-born neurons are actively traversing the intermediate zone, and P0, when migration is nearly completed, and neurons are positioned within the cortical plate (Fig 5B).

**Figure 5.**
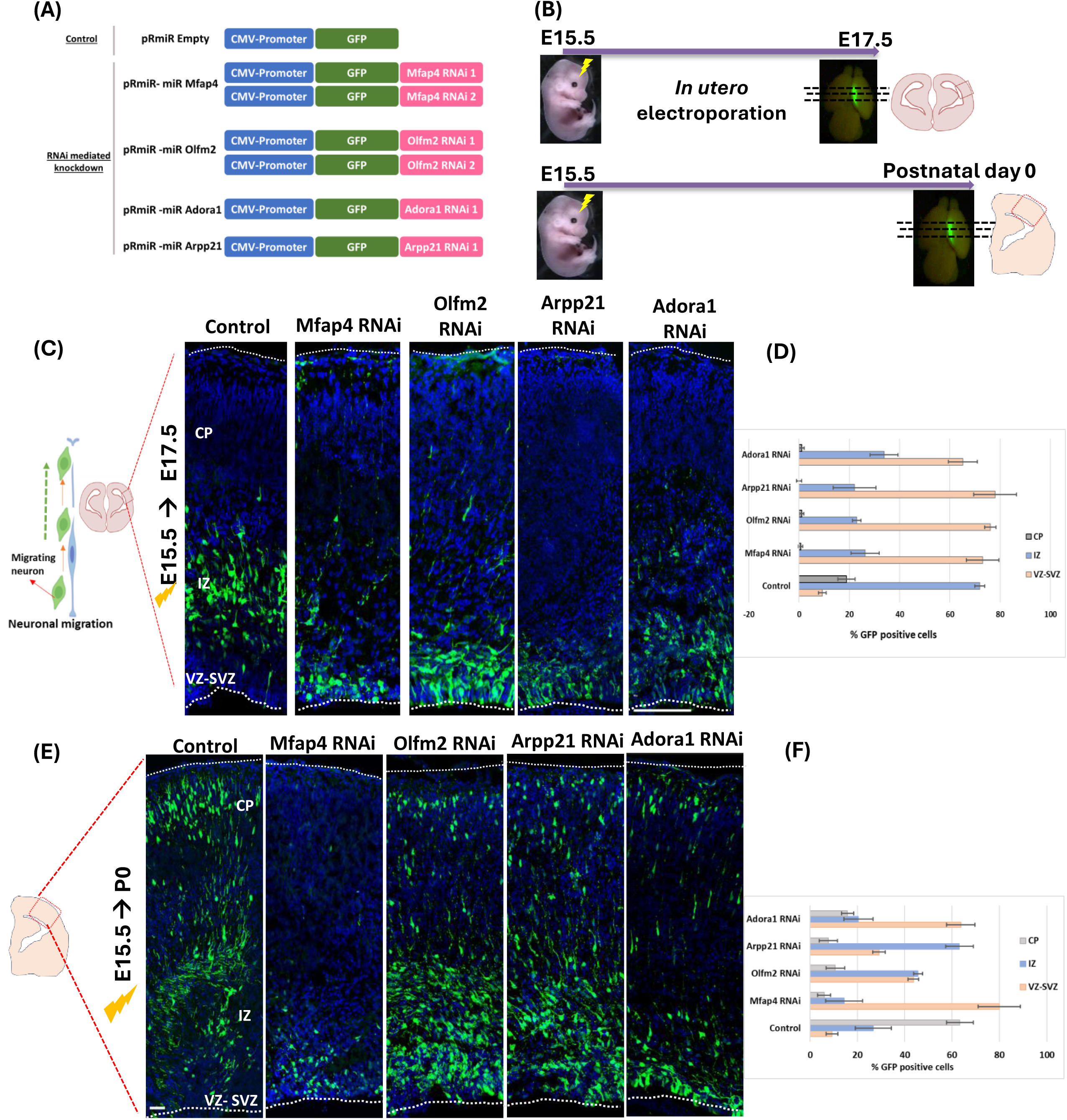
RNAi-mediated knockdown of BMP-responsive candidate genes disrupts radial migration of upper-layer cortical neurons. **(A)** Schematic of microRNA-based RNAi constructs used for gene-specific knockdown. miRNA sequences targeting the 3′ UTRs of candidate genes (*Mfap4, Olfm2, Adora1,* and *Arpp21*) were cloned into the pRmiR vector, which co-expresses GFP to label cells transfected through *in utero* electroporation. **(B)** Experimental design. Constructs were introduced into the embryonic mouse cortex by *in utero* electroporation at E15.5 to target progenitors generating upper-layer neurons. Brains were collected at E17.5 (active radial migration) and P0 (post-migratory laminar positioning) for analysis. **(C-D)** Representative coronal sections at E17.5 showing the distribution of GFP-positive neurons following control or gene-specific knockdown. In the control cortex, neurons have exited the ventricular zones and have populated the intermediate zone. Knockdown of *Mfap4* and *Olfm2* results in pronounced retention of neurons within the VZ–SVZ, with markedly reduced entry into the IZ and cortical plate. *Adora1* knockdown produces a substantial but less complete defect, while *Arpp21* knockdown shows strong accumulation of neurons in deeper zones and quantification of GFP positive cells across the cortex in control and RNAi electroporated E17.5 cortices (D). Scale bar: 50 µm. **(E-F)** Representative coronal sections at P0 showing final neuronal positioning. Knockdown of *Mfap4*, *Olfm2*, and *Arpp21* results in persistent mislocalization of neurons within the VZ–SVZ and IZ, indicating a severe migration defect. *Adora1* knockdown produces a partial phenotype, with a subset of neurons reaching the cortical plate, while the majority remain in deeper regions and quantification of GFP positive cells across the cortex in control and RNAi electroporated E17.5 cortices (F). Scale bar: 50 µm.

Knockdown of Mfap4 produced a pronounced arrest in neuronal migration when assessed 48 hours after electroporation at E17.5. In control cortices, the majority of GFP-positive neurons had exited the VZ-SVZ and occupied the intermediate zone, consistent with normal migratory progression at this stage (VZ-SVZ: 9.29 ± 1.54%; IZ: 71.88 ± 1.95%; CP: 18.83 ± 3.39%). Following Mfap4 knockdown, this distribution was sharply reversed: most GFP-positive cells remained confined to the VZ-SVZ (73.04 ± 6.42%), with few reaching the IZ (26.32 ± 5.61%) and essentially none entering the cortical plate (0.64 ± 0.89%). Olfm2 knockdown produced a comparable phenotype (Fig. 5C), with the majority of GFP-positive cells retained in the VZ-SVZ (76.05 ± 2.25%) and a few reaching the IZ and almost none to the CP (IZ: 22.90 ± 1.70%; CP: 1.04 ± 0.85%). Upon RNAi-mediated knockdown of Adora1 and Arpp21, we also observed significant migration defects with neurons failing to exit the proliferative zones within 48 hours of electroporation. The early migratory phase was disrupted for in both cases, with neurons accumulating in the VZ-SVZ at E17.5 (Adora1 RNAi: VZ-SVZ: [65.12± 5.73] %; IZ: [33.8 ± 5.47] %; CP: [1.07 ± 0.91] %; Arpp21 RNAi: VZ-SVZ: [77 ± 8.5] %; IZ: [22.08 ± 8.52] %; CP: [0.0 ± 0.0] %; Fig. 5C). These data indicate that these selected DEGs are required for the early steps of radial migration (Fig. 5C)

By P0, the effect of early arrest in migration was reflected in the continued mispositioning of GFP-positive neurons, with the majority remaining in the VZ–SVZ and IZ rather than reaching the cortical plate (Mfap4 RNAi: VZ-SVZ: [79.99 ± 8.88]%; IZ: [14.4 ± 7.87]%; CP: [5.94 ± 2.69]%; Olfm2 RNAi: VZ-SVZ: [43.67 ± 2.26]%; IZ: [45.62 ± 1.86]%; CP: [10.69 ± 3.94]%) (Fig. 5C). A similar mispositioning of GFP positive cells was observed following knockdown of Adora1 and Arpp21 (Fig. 5C-D). In control brains at P0, GFP-positive cells had largely completed their radial journey, with the majority distributed within the cortical plate (CP: 63.29 ± 5.61%; IZ: 26.69 ± 7.64%; VZ-SVZ: 9.25 ± 2.55%). Arpp21 knockdown produced a pronounced migratory block: the vast majority of GFP-positive cells were retained in the VZ–SVZ and IZ (VZ-SVZ: 29.12 ± 2.64%; IZ: 63.13 ± 5.81%), with few cells reaching the cortical plate (CP: 7.74 ± 3.80%). Adora1 knockdown yielded a milder but still pronounced defect: while a small fraction of neurons reached the cortical plate (CP: 15.83 ± 2.54%), the great majority remained in deeper compartments (VZ-SVZ: 63.74 ± 5.99%; IZ: 20.41 ± 6.16%), indicating a partial rather than complete migratory block.

For all four candidate genes, RNAi-mediated loss-of-function led to a significant retention of GFP-positive neurons in the lower region of developing cortices compared with controls across both time points. The severity of the phenotype varied among genes, ranging from the near-complete arrest produced by knockdown of Mfap4, Olfm2, and Arpp21 to the substantial but incomplete block observed with Adora1 knockdown; in each case, neurons failed to reach their normal laminar destination. The phenotypic convergence observed across four molecularly unrelated proteins reduces the likelihood of a shared nonspecific effect and is consistent with each gene representing a genuine downstream effector of BMP-regulated radial migration. The phenotype produced by each knockdown closely resembles the migration defect caused by dnBMPR1b-mediated inhibition of the BMP pathway at E15.5, in which neurons accumulate in the VZ-SVZ and intermediate zone rather than reaching the cortical plate (Saxena et al., 2018). Together, these results suggest that Mfap4, Olfm2, Adora1, and Arpp21 are functional downstream effectors of BMP signaling in regulating upper-layer cortical neuron migration. These findings also lend functional support for the transcriptional programs identified through spatial profiling and establish a mechanistic link between BMP-dependent gene regulation and the diverse cellular responses required for radial migration of upper-layer cortical neurons.

## Discussion

A central limitation in dissecting the mechanism through which extracellular signals regulate cortical neuronal migration has been the inability to link acute perturbation of a particular signaling pathway to spatially resolved, transcriptome-wide changes within a specific population of migrating neurons. Previous spatial transcriptomic studies of the developing cortex have relied primarily on unperturbed tissue, which has precluded linking the regulation of migration in distinct neuronal cohorts to any specific signaling pathway (Di Bella et al., 2021; Loo et al., 2019).

To address this gap, we paired temporally controlled *in utero* electroporation of a dominant-negative BMP receptor (dnBMPR1b), to perturb BMP signaling, with GeoMx Digital Spatial Profiling (DSP) at two developmental stages. This enabled the direct investigation of BMP-dependent transcriptional programs in E15.5-born upper-layer neurons during active migration (E17.5) and laminar positioning (P0), while preserving the spatial context. Our study reveals that BMP signaling engages largely non-overlapping transcriptional programs at these stages. Interestingly, we observed that the shared gene cohort between E17.5 and P0 was regulated in opposite directions upon perturbation of BMP signaling.

This pattern suggests that the transcriptional output of BMP signaling in post-mitotic migrating neurons is not static; rather, it is stage-dependent and is restructured as neurons transition from the intermediate zone to the cortical plate. Such context-dependence has been described for NOTCH signaling in progenitors (Kageyama et al., 2009). However, the BMP signaling-mediated temporal restructuring of transcriptional output in post-mitotic migrating neurons has not been investigated prior to this study.

We found that at E17.5, inhibition of BMP signaling leads to enrichment of chromatin-regulatory programs, including PRC2-mediated histone methylation, and reduction in *Actl6b*, a neuron-specific nBAF subunit encoding BAF53b (Lessard et al., 2007; Staahl et al., 2013). This suggests that BMP may contribute to maintaining nBAF-dependent chromatin accessibility as neurons traverse the intermediate zone. This interpretation is consistent with evidence that BAF complex disruption impairs radial migration through WNT-dependent transcriptional changes (Sokpor et al., 2021), since ACTL6B mutations cause epileptic encephalopathy and intellectual disability (Bell et al., 2019) and Wnt signaling has been shown to regulate neuronal migration and the multipolar to bipolar transition (Bocchi et al., 2017). At the progenitor level, stage-specific H3K27me3 patterning regulates the accessibility of genomic loci to BMP-activated SMADs (Katada et al., 2021). By analogy, a similar chromatin-based gating mechanism could operate in post-mitotic migrating neurons experiencing active BMP signaling, although this remains to be tested *in vivo*.

BMP inhibition at E17.5 is associated with a collective upregulation of ribosomal protein genes, a coordinated rank-level shift by GSEA (GOBP_CYTOPLASMIC_TRANSLATION, NES = +2.00, p = 0.0001) with only *Rpl12* crossing the individual fold-change threshold as well (Estimate = +0.54, p = 0.0012). This suggests that BMP signaling normally restrains translational capacity during active radial migration. However, this appears to reverse at P0, where 19 Rpl/Rps (ribosomal protein large/small subunit) genes are downregulated in dnBMPR1b neurons, indicating that at this stage BMP signaling supports ribosomal gene expression as neurons mature within the cortical plate. Since ribosomal protein composition influences the selective translation of specific mRNA subpools (Hornburg et al., 2014; Z. Shi et al., 2017), this stage-dependent reversal may reflect BMP-dependent modulation of translational state appropriate to each migratory phase, consistent with stage-dependent translatome remodeling in projection neurons (Glock et al., 2021; Shigeoka et al., 2016). One interpretation is that during the active migratory phase, neurons must engage substantial resources toward cytoskeletal remodeling and nuclear translocation; BMP-dependent restraint of ribosomal gene expression at this stage may reflect a prioritization of these motility demands over translational output. Conversely, as neurons settle into the cortical plate and begin elaborating dendrites and synaptic contacts, the translational burden increases sharply, and the shift toward BMP-supported ribosomal gene expression at P0 may reflect this transition from a motility program to a protein-synthesis-intensive maturation program.

At E17.5, elevated *Col4a5* and *Chpf2* expression in dnBMPR1b ROIs suggests increased collagen IV and chondroitin sulfate proteoglycan synthesis, respectively upon Bmp signaling inhibition. Since the CSPG-rich subplate/intermediate zone matrix is required for the multipolar-to-bipolar transition (Mencio et al., 2020; Mubuchi et al., 2024), elevated CSPG synthesis following BMP inhibition may perturb this transition, though this remains to be tested. *Adgrg1* (GPR56), which activates RhoA via Gα12/13 to inhibit neuronal migration(Luo et al., 2011; Singer et al., 2013), is also upregulated at E17.5; although GPR56-RhoA signaling normally operates at the pial surface rather than the intermediate zone, its ectopic upregulation here raises the possibility that ectopic activation of this pathway may contribute to migration arrest.

By P0, the dominant enriched programs shift toward synaptic specialization and membrane remodeling, with increased expression of synaptic adhesion molecules (*Lrfn5*, *Ptprd*), membrane-associated neuronal proteins (*Syngr1*, *Gpmcb*), and axon guidance receptors (*Rtn4r*, *Neo1*) in the dnBMPR1b ROIs. Since the ROIs at P0 were majorly placed within the cortical plate in both conditions, these transcriptional differences arise within neurons that have reached the similar anatomical compartment, consistent with a cell-intrinsic effect of BMP signaling on post-migratory transcriptional programs. Whether this reflects direct BMP-dependent transcription within CP-resident neurons or a difference in maturation state arising from the migration delay observed in dnBMPR1b neurons remains to be determined.

*Rtn4r* (*NgR1*), a GPI-anchored myelin receptor with established roles in restricting axon regeneration and synaptic plasticity in the mature cortex (McGee et al., 2005), is upregulated at P0 in neurons with dnBMPR1b expression. Upregulation of *Rtn4r* at this developmental stage before the onset of critical-period plasticity may reflect premature expression of circuit-stabilizing programs, although this interpretation requires direct testing. The upregulation of Rtn4r and Neo1 in dnBMPR1b expressing neurons located in the CP at P0 suggests that BMP-dependent transcriptional control extends to axon guidance and structural plasticity receptors as neurons consolidate laminar position. However, the functional consequences of their premature or elevated expression remain to be determined.

*Vldlr* (Very Low density lipoprotein receptor), which mediates the Reelin-dependent cortical plate stop signal (Hack et al., 2007; Trommsdorff et al., 1999), is also elevated, indicating that BMP signaling normally restrains neuronal responsiveness to the Reelin stop signal during laminar positioning. If this is true, then BMP inhibition-induced Vldlr upregulation could lower the Reelin concentration threshold required to arrest migration, potentially contributing to the laminar positioning defects observed in dnBMPR1b neurons, along with the failure to exit the IZ at earlier stages. Whether this reflects a direct transcriptional interaction, or a secondary consequence of altered maturation state cannot be resolved from the current data and would require further investigation.

Membrane lipid biosynthesis pathways enriched at P0 include genes such as *Smpd3* (nSMase2), encoding a key enzyme for sphingomyelin hydrolysis and ceramide generation in neurons (Stoffel et al., 2005), and *B3gnt5*, encoding a glycolipid glycosyltransferase contributing to sphingolipid and glycoprotein biosynthesis (Henion et al., 2001). *Gpat4*, which encodes a glycerophospholipid biosynthesis enzyme, additionally contributes to lipid pathway enrichment as a leading-edge gene across eight FDR-significant gene sets, although individually it narrowly misses the fold-change threshold (Est = +0.452; FDR = 0.025). This enrichment is consistent with the broad membrane remodeling that accompanies post-migratory neuronal maturation, albeit a direct link to BMP signaling remains to be established.

*Srf,* encoding the actin-dependent transcription factor, whose target program governs neuronal migration and hippocampal laminar organization (Alberti et al., 2005; Scandaglia et al., 2015; Stritt & Knöll, 2010), shows a nominal decrease at P0 (Estimate = -0.487; FDR = 0.013). This is consistent with *Srf* appearing as a leading-edge contributor to the negatively enriched actomyosin structure organization gene set at P0 (NES = -1.61), but narrowly misses the |log₂FC| > 0.5-fold-change gate applied throughout this study. As a near-threshold candidate with strong statistical support, it raises the possibility that BMP signaling sustains SRF-dependent actin cytoskeletal programs during post-migratory maturation. Similarly, *Maml3*, a Notch transcriptional coactivator with additional nuclear coactivator functions (Alves-Guerra et al., 2007; Jin et al., 2010; Oyama et al., 2011; Zhao et al., 2007), shows opposing stage-dependent regulation; being upregulated at E17.5 (Est = +0.715) and downregulated at P0 (Est = -1.006). This places *Maml3* among the 17 genes exhibiting directional reversal at the stage overlap. However, whether its stage-dependent regulation reflects direct BMP-SMAD control of Notch coactivator activity, remains to be determined.

Knockdown of four molecularly unrelated candidates-an ECM glycoprotein (Mfap4), a secreted olfactomedin (Olfm2), a G-protein-coupled receptor (Adora1), and an RNA-binding regulator (Arpp21); each produced a similar phenotype of migratory arrest, indicating that BMP coordinates migration through parallel transcriptional outputs rather than a single effector pathway.

Mfap4/MFAP4 is a fibrillin-interacting ECM glycoprotein that engages integrin αvβ3 via an RGD motif to promote cell migration and matrix organization (Wozny et al., 2024). MFAP4 protein occupies meningeal and perivascular compartments in the human CNS (Samadzadeh et al., 2023). Expression analysis conducted through mRNA *in situ* hybridization in this study shows that the *Mfap4* transcript is primarily present in the lower region of the developing cortex at E15.5, at the site of the multipolar-to-bipolar transition and initiation of locomotion of newborn neurons. Furthermore, we found that BMP signaling exerts opposing effects on Mfap4 across stages, since inhibition of BMP signaling elevates expression at E17.5 (Estimate = +1.44, p = 0.008, FDR = 0.56) and reduces it at P0 (Estimate = -0.74, p = 0.036, FDR = 0.37), suggesting that while BMP normally restrains the expression of Mfap4 during IZ transit, it switches to increasing Mfap4 expression during later stages of migration. This reversal is consistent with stage-dependent modulation of ECM-related processes, potentially influencing integrin-mediated adhesion during migration and subsequent laminar organization. In fact, we observed that knockdown of Mfap4 at E15.5 produces a near-complete migration block by E17.5, which continues till P0, establishing its functional requirement across the active migratory phases.

Olfm2, a secreted olfactomedin-domain protein enriched in the lower region of the developing mouse cortex (Sultana et al., 2014), is downregulated upon BMP inhibition at E17.5 (Estimate = - 1.34, p = 0.026, FDR = 0.63). Olfm1/Noelin1 is known to stabilize surface AMPA receptor complexes (Pandya et al., 2018); similarly, Olfm2 has been shown to associate with the GluR2 subunit of AMPAR complexes in the cortex (Sultana et al., 2014), suggesting a shared role in cell-surface receptor stabilization. Notably, Olfm2 is also transcriptionally induced downstream of TGF-β superfamily signaling and mediates cytoskeletal gene expression through SRF (N. Shi et al., 2014), raising the possibility that its BMP-dependent regulation in cortical neurons similarly converges on actin cytoskeleton dynamics relevant to migration; however, this connection remains speculative. At the GSEA level, Olfm2 contributes to the negatively enriched synaptic membrane compartment at E17.5 (NES = -1.40) and the same gene set in its positively enriched form at P0 (NES = +2.22), mirroring the stage-dependent transcriptional reversal; its expression also trends upward in dnBMPR1b ROIs at P0 (Estimate = +0.457, p = 0.106), though this does not reach significance.

The reduction of *Adora1* in dnBMPR1b ROIs at E17.5 (Estimate = -0.826, p = 0.012, FDR = 0.58) places purinergic signaling within the BMP-dependent transcriptional program during active radial migration. At the GSEA level, Adora1 is a leading-edge gene in the negatively enriched synaptic physiology-related sets at E17.5 and the positively enriched synaptic membrane sets at P0. A1R normally suppresses cAMP/PKA signaling via Gᵢ (Fredholm et al., 2011), and PKA regulates centrosomal dynamics during nucleokinesis in cortical interneurons (Stoufflet et al., 2020); whether this mechanism also operates in glial-guided excitatory neurons remains to be determined. It is possible that BMP-dependent expression of Adora1 during IZ transit may tune cAMP tone at a stage when cytoskeletal dynamics are critical.

Arpp21 is reduced in dnBMPR1b expressing neurons (Estimate = -1.099, p = 0.017, FDR = 0.58). ARPP21 normally buffers miR-128 activity by binding and antagonizing it, thereby stabilizing its neurodevelopmental mRNA targets (Rehfeld et al., 2018). Premature miR-128 expression alone is sufficient to impair upper-layer neuron radial migration (Franzoni et al., 2015). BMP inhibition would therefore be expected to lower ARPP21 levels, release miR-128 from antagonism, and permit degradation of migration-relevant transcripts; a possible mechanism supported by the migration defect produced by Arpp21 knockdown.

There are some limitations of the current study that warrant consideration. For example, GeoMx DSP profiles population-average transcriptomes from manually delineated ROIs, such that the observed changes may reflect contributions from neurons at slightly different stages of differentiation or from residual progenitors, though cell-type deconvolution confirms migrating neurons as the predominant cell type in E17.5 GFP-positive ROIs (mean abundance scores of 103 and 111 for control and dnBMPR1b conditions).

The stage-resolved DEG datasets generated through this study provide an entry point for assembling the BMP-responsive gene regulatory network and assigning regulatory hierarchy among its components. In fact, the approach combining temporally controlled pathway perturbation with spatial transcriptomics in defined neuron populations offers a scalable framework for dissecting the transcriptional logic underlying cortical lamination in other signaling contexts. Importantly, the data obtained through this study establishes that BMP signaling is not a static pathway with a fixed transcriptional output. The pathway’s targets, their direction of regulation, and the cellular processes they serve are all subject to temporal reorganization as the neuronal context shifts from active migration to laminar positioning. This stage-dependence of signaling output may be a general principle by which developmental morphogens operating through the same receptor and downstream effector cascades achieve context-appropriate transcriptional responses. In the case of BMP signaling, whether this requires parallel upstream diversification or emerges from cellular-context-dependent interpretation of a shared SMAD cascade remains to be determined. The mechanistic contribution of this study has been the demonstration of this same principle in the specific context of radial migration of E15.5-born cortical neurons, and has resulted in the identification of four effector genes that bridge pathway activation to cellular behavior.

## MATERIALS AND METHODS

### NanoString GeoMx DSP digital spatial transcriptomics and sequencing: Slide Preparation

Control and test (dnBMPR1b)-electroporated brains were harvested at E17.5 and P0, processed and embedded following a modified manufacturer’s protocol, and sectioned coronally into 8–10 µm cryosections. Slides were stored at −80 °C until further processing. For GeoMx analysis, slides were thawed, fixed overnight in 10% neutral-buffered formalin (NBF) at room temperature, and washed with PBS. Pre-baking imaging was performed to preserve GFP fluorescence, as recommended in the GeoMx workflow. Slides were then ethanol-dehydrated, baked at 65 °C, subjected to high-pH antigen retrieval using Invitrogen IHC Antigen Retrieval Solution (Cat#00-4956-58) according to the manufacturer’s instructions (Thermo Fisher Scientific, 2020). Protease digestion, post-fixation, and NBF stop-buffer washes were performed as described in the GeoMx RNA-NGS Slide Preparation Manual (NanoString Technologies, 2023b).

Slides were hybridized overnight at 37 °C with GeoMx Mouse Whole Transcriptome Atlas (WTA) probes (NanoString Technologies, 2023a), followed by stringent formamide/SSC washes. After blocking with Buffer W, tissues were stained with SYTO 13 nuclear dye and mounted on the GeoMx Digital Spatial Profiler for imaging and ROI selection (NanoString Technologies, 2023b).

### ROI selection and collection of indexing oligonucleotides from ROIs

The slide was imaged at 20X magnification, and ROIs were selected using the overlay feature of GeoMx DSP. A JPEG file of the same slide previously imaged was overlaid on the new scan, and Polygonal ROIs were placed based on the GFP signal of control or dnBMPR1b electroporated mouse cortices. A total of 23 ROIs were selected from E17.5 and P0 sections. With a microcapillary, the indexing oligonucleotides from the regions of interest were released by exposure to 385 nm light (UV) and deposited into a 96-well plate (Danaher et al., 2022; NanoString Technologies, 2023a).

### Library preparation and sequencing

The 96-well plate was dried in a PCR machine at 65 °C for 1 hour and then reconstituted in 10 μL of DEPC-treated water (Ambion, AM9922). Sequencing libraries were prepared according to the NanoString GeoMx-NGS Readout Library Prep manual (Griswold, Matthew et al., 2025; NanoString Technologies, 2023b; Shaikh et al., 2024). The PCR reaction was performed using NanoString SeqCode primers. A total of 21 PCR cycles were performed according to the conditions outlined in the manual. PCR products were then pooled in equal volumes and purified using AMPure XP beads (Beckman Coulter, A63880) twice. The library was sequenced on an Illumina NovaSeq 6000, using 2x 151 bp paired-end reads.

### NanoString GeoMx WTA data analysis

The sequenced FastQ files were further processed to convert them into dcc (Digital Count Conversion) format using the GeoMx NGS Pipeline v.2.3.3.10 from NanoString. The trimming of adapters, removal of duplicates, and mapping to the reference probe barcodes present in the WTA panel, as well as quantification, were performed during the conversion process from fastq to dcc files. The individual ROI thus results in a single .dcc file The resulting dcc files from each ROI were further imported and analyzed using the GeoMxTools R package (Griswold, Matthew et al., 2025). R version 4.2 (https://www.r-project.org/about.html) was used throughout the analysis unless stated otherwise. The analysis within the GeoMxTools pipeline includes QC for both Probe and Segment, Normalization, and Unsupervised clustering. The parameters used during the QC of segments includes: minSegmentReads = 1000, percentTrimmed = 80, percentStitched = 80, percentAligned = 80, percentSaturation = 50, minNegativeCount = 10, maxNTCCount = 1000, minNuclei = 200, minArea=5000 and minLOQ = 2. Due to the inherently sporadic and mosaic nature of *in utero* electroporation, few ROIs did not reach the recommended minimum nuclei threshold but were retained in this study. The parameters used for the QC or filtering of probes (target genes) include: geometric mean of each probe’s counts from all segments divided by the geometric mean of all probe counts representing the target from all segments < 0.1, and percentage of segments within which the probe is identified as an outlier based on Grubb’s test >= 20%. A probe is removed locally (from a given segment) if it is an outlier according to Grubb’s test in that segment (LOQ: Limit of Quantification).

After appropriate QC on segments and target genes, the counts were normalized using the Q3 method of normalization. The differential gene expression testing was performed using the LMM (Linear Mixed Models) within the GeoMxTools R package, which adjusts for the fact that multiple regions of interest are placed per tissue section and are not independent observations, as understood by other statistical tests. Genes with a P-value < 0.05 were considered statistically significant, with an Effect size > 0.5 indicating upregulation and < -0.5 indicating downregulation.

For the deconvolution of selected cell type abundance within the ROIs, the SpatialDecon algorithm (Danaher et al., 2022) was used, where a custom gene expression profile matrix was generated from single-cell RNA-Seq data of GSE123335 for the P0 stage and GSE153162 for the E17.5 stage ROIs. SpatialOmicsOverlay R-package (https://www.bioconductor.org/packages/release/bioc/html/SpatialOmicsOverlay.html) was utilized for overlaying the expression of selected markers over the GeoMx scan image within selected ROIs (Griswold, Matthew & Danaher, Patrick, 2023).

### Functional Gene Set Enrichment Analysis

Gene set enrichment analysis (GSEA) was performed to evaluate the upregulated and downregulated Reactome pathways, as well as gene ontology categories (Biological Process, Cellular Component, and Molecular Function), within each comparison using the clusterProfiler R package (Yu et al., 2012). The individual gene sets for the GSEA test were obtained from the MSigDB database (Liberzon et al., 2015). Default settings were used with 1000 phenotype permutations to generate the P-value and FDR values. Gene sets were considered significantly different between the compared groups with P-value < 0.05.

### Transcription factor binding site screening

Predicted transcription factor (TF) binding motifs were analyzed using the JASPAR TFBS extraction tool (Fornes et al., 2020) (https://jaspar.elixir.no/tfbs_extraction/) within the 2-kb promoter region and an additional 100 bp downstream of target mouse genes (using JASPAR2024), prioritizing motifs with a JASPAR score > 400. In addition, HOMER (Hypergeometric Optimization of Motif EnRichment) (Heinz et al., 2010) was utilized for both known and novel motif enrichment analysis, using the script findMotifs.pl and setting parameters specific to the mouse. HOMER predicted the motif with the lowest p-value as the key transcription factor.

### Identification of BMP-responsive elements

Promoter regions of well-known BMP target genes are known to possess GC-rich motifs (GGCGCC), also known as BMP-responsive elements (BRE). The GC-rich motif was screened within the promoter of target genes using FIMO from The MEME Suite web server (https://meme-suite.org/meme/doc/fimo.html).

### Statistical Analysis

R Statistical Software was used for all statistical analyses (version 4.2.1; R Foundation for Statistical Computing, Vienna, Austria). Differential expression of individual genes was performed using a linear mixed model (LMM) method, with the assumption that several ROIs were taken from the same tissue (random effect). To account for multiple comparisons, we adjusted the resulting p-values with the FDR procedure to control the false discovery rate. We used the Kruskal–Wallis test to examine differences in gene expression across the ROI groups, followed by pairwise Wilcoxon tests and statistical significance was defined as a p-value < 0.05.

### Experimental animals

All animal experiments were performed according to the protocol IITK/IAEC/2020/1118 approved by the Institute Animal Ethics Committee, registered with CPCSEA (Reg. no. 810/ 03/ ac/ CPCSEA, dated 15/10/2003). Males and female wild-type mice (C57BL/6JNcbs, Stock no.000664; Jackson Laboratory) were crossed to generate timed pregnant females for the in-utero electroporation experiments and embryonic or postnatal brain harvesting.

### In utero electroporation

Timed pregnant females (E15.5) were anesthetized using an isoflurane vaporizer (Patterson Veterinary). Uterine horns of the pregnant female were exposed and the purified plasmid DNA construct(s) at a total concentration of 1 μg/μl along with 0.05% (w/v) Fast Green FCF was injected into the forebrain vesicle (lateral ventricle) of the embryos through the uterine wall into the lateral ventricle of the embryos using a glass microcapillary (Sutter Instruments-BF150-86-10) and aspirator tube assembly *(Sigma-A5177-5EA).* Electroporation was performed with tweezer-disc electrodes (CUY650-10, Nepagene) providing an electric pulse of 35 V for 50 ms, five times at an interval of 950 ms, using an electroporator (ECM 830, BTX) (Saxena et al., 2018).

### Constructs

For spatial transcriptomic profiling: pCAG-dnBMPR1b-IRES-GFP, pCAG-IRES-GFP were used for *in utero* electroporation. For functional validation of DEGs: pRmiR-Mfap4-RNAi1, pRmiR-Mfap4-RNAi2, pRmiR-Olfm2-RNAi1, pRmiR-Olfm2-RNAi2, pRmiR-Arpp21-RNAi, pRmiR-Adora1-RNAi, and pRmiR-Empty vector (Udaykumar et al., 2023) were used.

### Cloning of constructs

#### Cloning of pCAG-dnBMPR1b-IRES-GFP

The dnBMPR1b coding sequence from pCAG-dnBMPR1b (Saxena et al., 2018) was subcloned between the EcoRI and NotI sites in pCAG-NeuroD1-IRES-GFP (Addgene #45025), by replacing the NeuroD1 coding sequence, giving pCAG-dnBMPR1b-IRES-GFP.

#### Designing and cloning of miRNA Oligos

To achieve knockdown of selected DEGs, microRNAs (miRNAs) targeting the 3’ untranslated region (UTR) of mouse Mfap4, Olfm2, Adora1, and Arpp21 were designed utilizing the BLOCK-iT RNAi Designer (Invitrogen). The sequences of the oligos are given in the table. The synthesized miRNA oligonucleotides, top and bottom strands, were annealed, and the resulting duplexes were ligated into the BsaI-digested pRmiR vector (Smith et al., 2009).

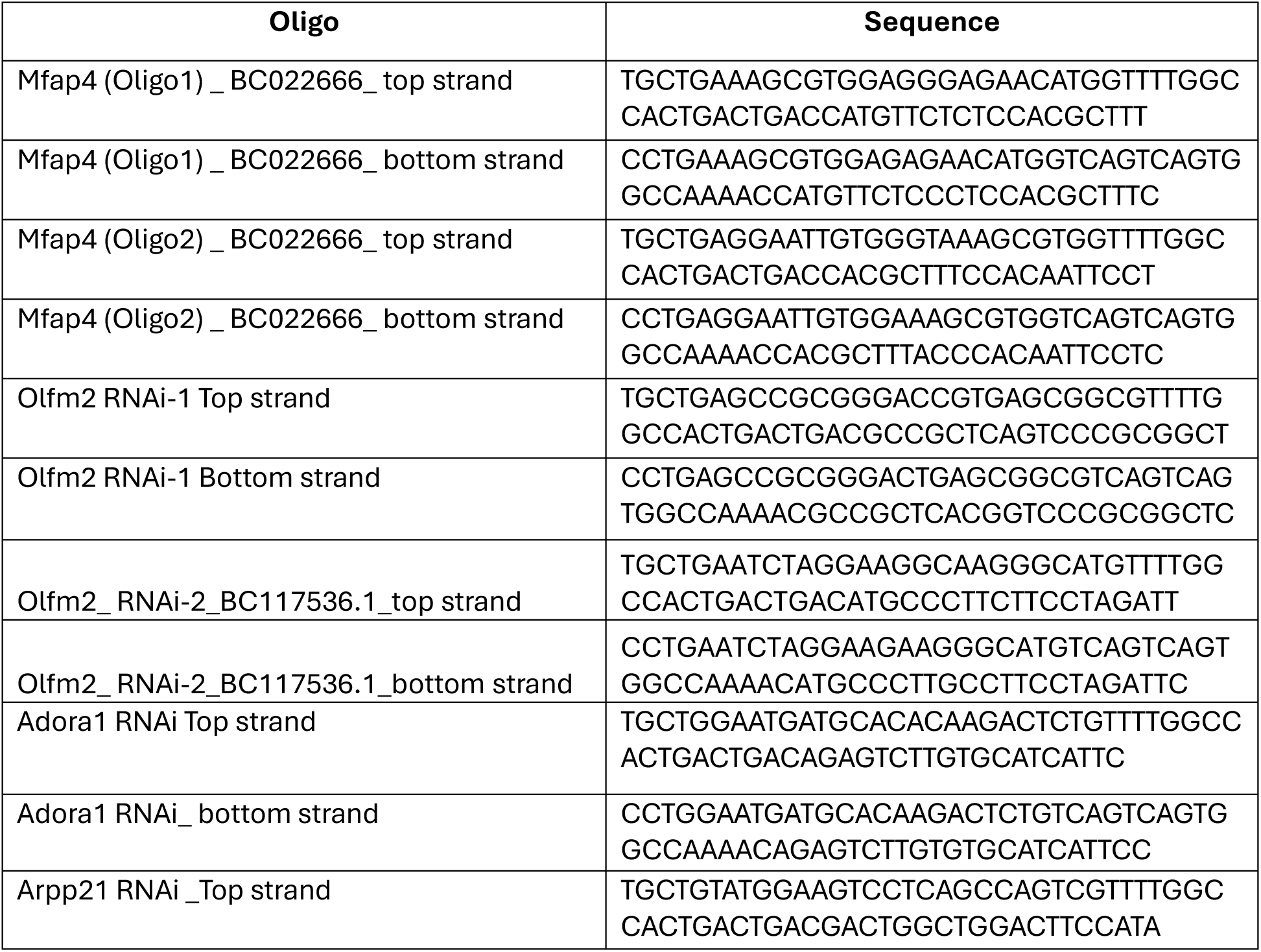
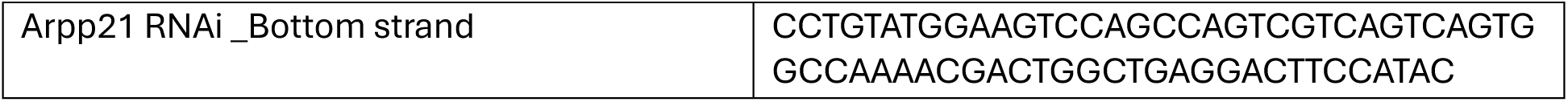

### Immunohistochemistry

Cryosections were thawed and air-dried before being rinsed with 1× PBS and fixed in 4% PFA for 10 min. The sections were then permeabilized using 0.1% PBT (1× PBS containing 0.1% Tween-20) and blocked for 1 h at room temperature in 5% heat-inactivated goat serum (HINGS) prepared in PBT. Following blocking, the sections were incubated overnight at 4 °C with using anti-GFP (1:500; A-6455, ThermoFisher Scientific). This was followed by incubation with the corresponding secondary antibodies at room temperature. The sections were counterstained with DAPI (Sigma-Aldrich, D9542), followed by three washes with PBT for 5 min each. Finally, the sections were mounted using an antifade mounting medium.

### RNA *in situ* hybridization

RNA *in situ* hybridization was performed on the coronal forebrain sections of unmanipulated or in utero electroporated C57BL/6JNcbs mice according to established protocols (Trimarchi et al., 2007). RNA *in situ* hybridization experiments were performed independently three times (n=3 independent biological replicates) to confirm reproducibility of the observed expression patterns. The digoxigenin-labeled antisense riboprobes for RNA *in situ* hybridization were made through in vitro transcription using the following mouse cDNA clones (Mouse full-length cDNA library, Thermofisher, transOMIC technologies): Mfap4 (Accession number: BC022666), Spag5 (Accession number: BC052672), Zfp219 (Accession number: BC071271), Zfp692 (Accession number: BC150776), Zfp3 (Accession number: BC096463), Pcdhb22 (Accession number: BC094239), Olfm2 (Accession number: BC117536), Adora1 (Accession number: BC079624), Arpp21 (Accession number: BC053001).

### Image acquisition and processing

Fluorescence images of immuno-stained electroporated mouse cortical sections were acquired using an AXR Nikon confocal microscope and processed with NIS Elements AR software. Images from RNA *in situ* hybridization experiments were captured using a Leica DM5000B automated upright microscope equipped with a DFC7000T camera and Leica Application Suite X (LAS X) software. Representative images of electroporated mouse brains were acquired using a Leica MZ10F stereomicroscope fitted with a DFC420C camera. Basic image processing was performed using Adobe Photoshop 26.11.3. GFP-positive cells in the electroporated cortical sections were manually quantified using the Count Tool in Photoshop. Scale bars were measured using the built-in Ruler Tool in Adobe Photoshop, with standard pixel-size values corresponding to the respective magnifications.

### Quantification and statistical analysis

For quantification of GFP-positive cells in electroporated mouse cortices, each electroporated embryo was considered an individual biological replicate (n). For each biological replicate, three cortical sections were analyzed as technical replicates. GFP-positive cells from all three technical replicates were counted and pooled for each biological replicate, and the mean value was calculated (N). The mean and standard deviation (S.D.) presented in the graphs were calculated from the values obtained across all biological replicates. Statistical analyses were performed using GraphPad Prism 8.0.2, and the data are presented as mean ± S.D. (%) of GFP-positive cells. Statistical significance was assessed using an unpaired Student’s t-test.

## Footnotes

### Author contributions

N.A., and J.S., designed research; N.A., A.J., M.M., V, B., performed research; N.A., A.J., and J.S. analyzed data; and N.A., A.J., M.M., and J.S. wrote and reviewed the manuscript and J.S. supervised the project, administered the project, and acquired funding.

### Declaration of generative AI and AI-assisted technologies

During the preparation of this manuscript, the authors used OpenAI and Claude to assist with improving the language, clarity, readability, and phrasing in certain sections. The authors reviewed and edited those sections as appropriate and take full responsibility for the accuracy, integrity, and final content of the published work.

## Acknowledgments

The authors acknowledge use of the confocal facility supported by ICMR-CoE grant 5/3/8/3/2021/ITR to Prof. Amitabha Bandyopadhyay and Prof. S. Ganesh. The authors also thank Mr. Naresh Gupta for his technical support with this study.

## Supplementary files

Supplementary file and additional raw data files (.xlsx) files are provided.

## Data availability

All data supporting the conclusions can be found in the main text or are available upon request from the lead contact (N.A.), Any additional information required to reanalyze the data reported in this study is available from the lead contact upon request. Requests for further information and resources should be directed to and will be fulfilled by the lead contact (N.A.), corresponding author (J.S.).

