## Supplementary material for "Spatial transcriptomics reveals BMP-dependent stage-specific transcriptional programs underlying migration of cortical neurons": Supplemenary file

**Supplementary Fig. S1**

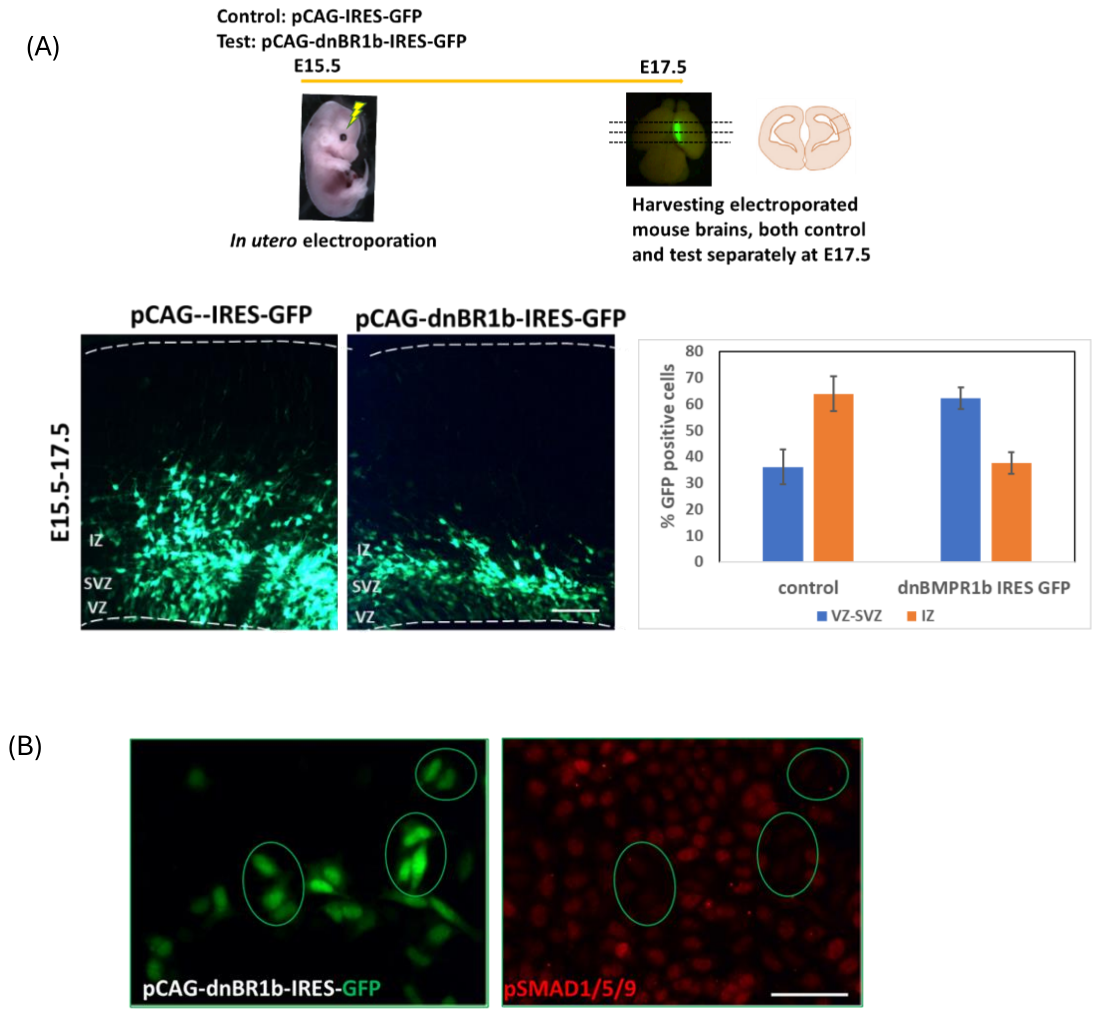


(A) Schematic of experimental timeline and E17.5 cortex electroporated with control plasmid and dnBMPR1b-IRES-GFP depicting distribution of GFP positive cells across cortex (control: VZ-SVZ: [36 ± 6.6] %; IZ: [64 ± 6.60] %; dnBMR1b IRES GFP: VZ-SVZ: [62.3 ± 4.09] %; IZ: [37.7 ± 4.1] %). Data are presented as mean ± s.d; unpaired student t-test. Differences were considered statistically significant at $*p < 0.05$. (VZ: Ventricular zone, SVZ: Subventricular zone, IZ: Intermediate zone. Scale bar:100 µm. (B) Immunocytochemistry of pSMAD1/5/9 on HEK 293T cells, which have been transfected with pCAG-dnBMPR1b-IRES-GFP (Scale bar:100 µm).

**Supplementary Fig. S2**

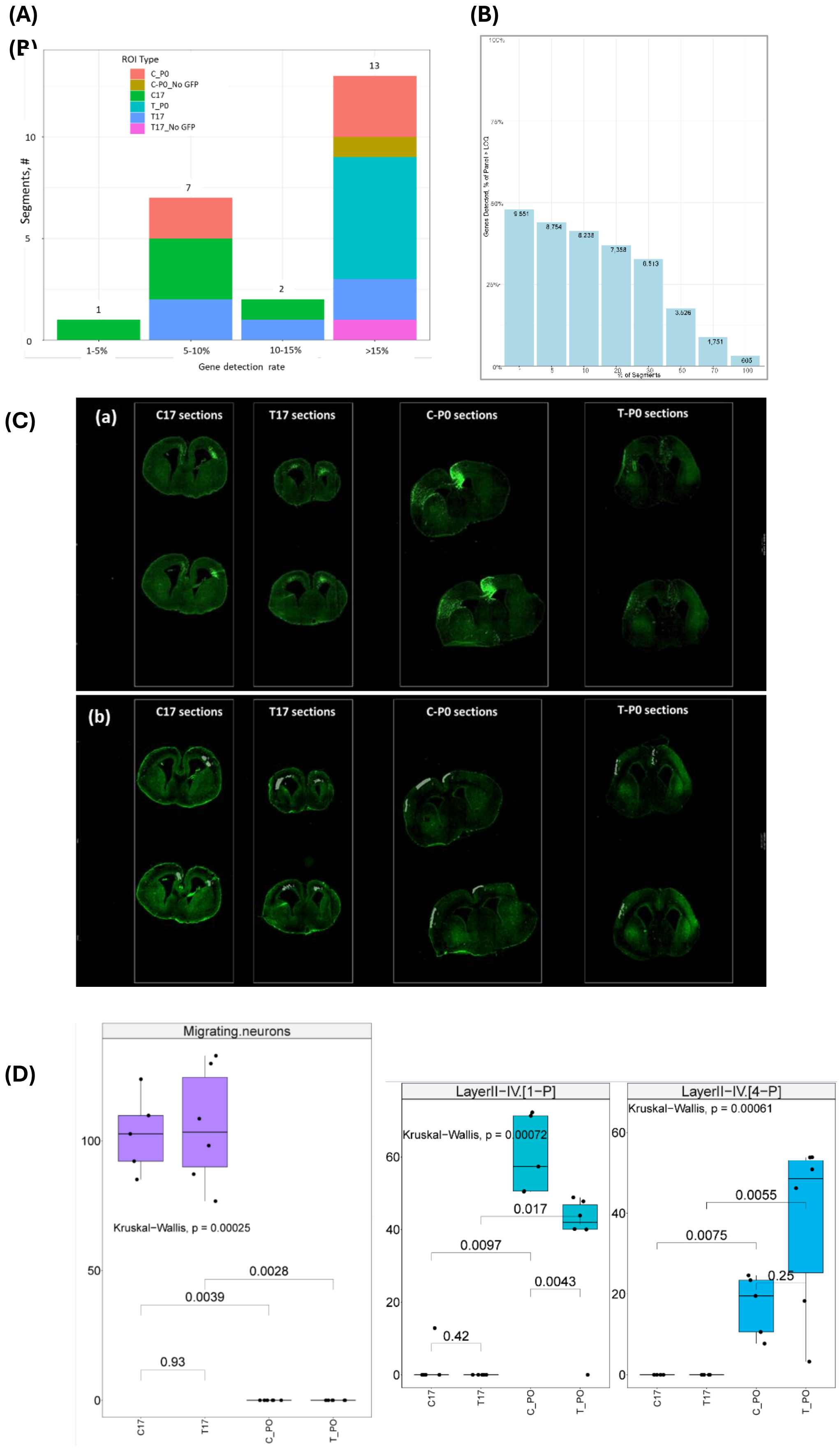

(A) Gene detection rate per ROI type, (B) Gene detection across segments, (C) Whole GeoMx slide image showing all the electroporated mouse brain sections from C17, T17, and C-P0, T-P0 (a), along with corresponding ROIs marked onto these sections (b) that were used in the study, (D) Cell-type abundance within the selected ROIs was estimated using the SpatialDecon algorithm with custom gene expression profile matrices generated from stage-matched single-cell RNA-seq datasets from GSE153162 for E17.5 and GSE123335 for P0. The reference profiles were derived from the respective published single-cell transcriptomic studies (Loo et al., 2019; Di Bella et al., 2021). Box plots show cell-type abundance scores for individual ROIs, with the cell populations represented as indicated in the plots. Statistical comparisons were performed using the Kruskal-Wallis test, with P values indicated in the plots.

**Supplementary Fig S3: RNA *in situ* hybridization-based expression screening of selected DEGs.**

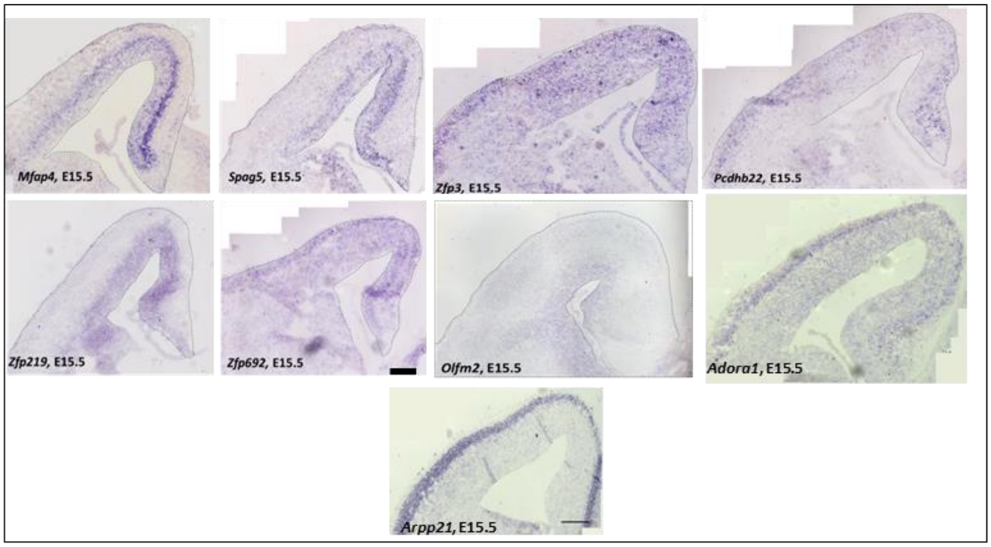


RNA *in situ* hybridization-based expression screening of Mfap4, Spag5, Zfp3, Pcdhb22, Zfp219, Zfp692, Olfm2, Adora1, Arpp21 performed on WT E15.5 mouse cortices (Scale:100 μm).
